# The recycling endosome protein Rab25 coordinates collective cell movements in the zebrafish surface epithelium

**DOI:** 10.1101/2021.01.29.428759

**Authors:** Patrick Morley Willoughby, Molly Allen, Jessica Yu, Roman Korytnikov, Tianhui Chen, Yupeng Liu, Isis So, Neil Macpherson, Jennifer Mitchell, Rodrigo Fernandez-Gonzalez, Ashley E. E. Bruce

**Affiliations:** Cell and Systems Biology, University of Toronto, Toronto, ON, Canada; Ted Rogers Centre for Heart Research, Translational Biology and Engineering Program, University of Toronto, ON, Canada; Institute of Biomaterials and Biomedical Engineering, University of Toronto, Toronto, ON, Canada

**Keywords:** zebrafish, epiboly, epithelial morphogenesis, Rabs, trafficking, cytokinesis, abscission, viscoelasticity

## Abstract

In emerging epithelial tissues, cells undergo dramatic rearrangements to promote tissue shape changes. Dividing cells remain interconnected via transient cytokinetic bridges. Bridges are cleaved during abscission and currently, the consequences of disrupting abscission in developing epithelia are not well understood. We show that the Rab GTPase, Rab25, localizes near cytokinetic midbodies and likely coordinates abscission through endomembrane trafficking in the epithelium of the zebrafish gastrula during epiboly. In maternal-zygotic Rab25a and Rab25b mutant embryos, morphogenic activity tears open persistent apical cytokinetic bridges that failed to undergo timely abscission. Cytokinesis defects result in anisotropic cell morphologies that are associated with a reduction of contractile actomyosin networks. This slows cell rearrangements and alters the viscoelastic responses of the tissue, all of which likely contribute to delayed epiboly. We present a model in which Rab25 trafficking coordinates cytokinetic bridge abscission and cortical actin density, impacting local cell shape changes and tissue-scale forces.

## Introduction

During metazoan development, epithelial cells are often organized into cohesive sheets that undergo large-scale cellular movements. In proliferative tissues, cell division is fundamental to the establishment of normal tissue architecture (Gibson et al., 2006). Cell division also poses a challenge for developing epithelia, as dividing cells must maintain cell-cell contacts while simultaneously coordinating shape changes with surrounding cells (Higashi et al., 2016). In the zebrafish embryo, the single-cell thick surface epithelium, the enveloping layer (EVL), undergoes rounds of proliferation during epiboly, a gastrulation movement describing the expansion and spreading of a tissue (Campinho et al., 2013). Thus, the zebrafish gastrula is an ideal system to investigate epithelial morphogenesis during gastrulation.

The multilayered zebrafish blastoderm is positioned on top of a large yolk cell. During epiboly, the blastoderm spreads vegetally to completely cover the yolk cell (reviewed in Bruce and Heisenberg 2020). Epiboly initiates at dome stage when EVL spreading is triggered by a reduction in tissue surface tension, that enables vegetal directed tissue expansion (Morita et al., 2017). EVL cells thin apicobasally as the apical surface area expands, with cell divisions preferentially oriented along the animal-vegetal axis (Campinho et al., 2013). EVL proliferation slows as the blastoderm covers half of the yolk cell at 50% epiboly (Campinho et al., 2013).

At 6hpf, a contractile actomyosin ring assembles in the region of the yolk cell adjacent to the blastoderm, the yolk syncytial layer (YSL) (Berhndt et al., 2012). Contractile forces generated by the ring tow the tightly attached EVL vegetally. During the later stages of epiboly, as EVL cells continue to elongate and spread, EVL tension is highest, particularly in marginal regions closest to the yolk cell (Campinho et al., 2013). A subset of marginal EVL cells actively change shape and rearrange to leave the margin, enabling the tissue circumference to narrow and eventually close at the vegetal pole (Köppen et al., 2006). Overall, EVL morphogenesis involves coordination between local cell shape changes, rearrangements and tissue-scale forces.

Cell divisions play a critical and conserved role during epithelial morphogenesis in multiple systems (Campinho et al., 2013; Firmino et al., 2016; Lau et al., 2015; Wyngaarden et al., 2010). Following division, daughter cells either share a common interface or cell rearrangements occur (Firmino et al., 2016; Gibson et al., 2006). Both events can be viewed as multicellular processes as cell division impacts both the dividing cell and adjacent neighbours. For example, during the formation of *de novo* cell contacts between newly formed daughter cells, neighbouring cells resist contractile forces to maintain shape as shown in the Xenopus gastrula and Drosophila pupal notum (Herszterg et al., 2013; Higashi et al., 2016).

Cell divisions also impact tissue scale forces. Oriented cell divisions have well characterized roles in both propagating forces and dissipating anisotropic tensions (reviewed in Godard and Heisenberg, 2019). During zebrafish epiboly, oriented cell divisions relieve stresses in the EVL to facilitate tissue spreading (Campinho et al., 2013). More recently, cell divisions have been shown to contribute to the viscoelastic properties of cells. For example, the deep cells in the central region of the zebrafish blastoderm undergo a rapid decrease in tissue viscosity following mitotic rounding that is important for initiating epiboly (Petridou et al., 2019a). Similarly, inhibiting cell division in amniote gastrulation resulted in high tissue viscosity, slowing cell Polonaise movements (Saadaoui et al., 2020). While our understanding of the effects of cell divisions on tissue development has advanced considerably, the role of the terminal steps of cytokinesis, intercellular bridge abscission has not been examined in detail.

Following contraction of the cytokinetic ring, sister cells remain connected via intercellular bridges (Burgos and Fawcett, 1955). Actomyosin and microtubule networks constrict the bridges to 1-2μM (Mierzwa and Gerlich, 2014) and subsequently, both cytoskeletal networks are removed from bridges which is followed by bridge cleavage adjacent to the cytokinetic midbody (Connell et al., 2009; Frémont et al., 2017; Mierzwa and Gerlich, 2014). The timing of bridge abscission can affect many biological processes. When abscission is delayed in cancer cell culture models, neighbouring cell division events can tear open adjacent cytokinetic bridges resulting in the formation of binucleate cells (Dambournet et al., 2011). In mouse ES cells, abscission impacts cell fate by acting as a switch for pluripotency exit (Chaigne et al., 2020). Lastly, cytokinetic bridges act as landmarks for lumen development in both cell culture and zebrafish, with the timing of abscission critical for successful lumen expansion (Rathbun et al., 2020; Willenborg and Prekeris, 2011). More specifically, in Kupffer’s vesicle, the zebrafish leftright laterality organ, premature cytokinetic bridge cutting via laser ablation disrupted lumen morphogenesis (Rathbun et al., 2020). Open questions include what happens to cells if they remain interconnected by cytokinetic bridges during tissue morphogenesis and do cytokinesis failures hinder cell rearrangements or disrupt tissue-scale forces? Furthermore, what subcellular pathways are required for abscission during embryonic development?

Rab proteins are the largest family of small GTPases and are important for intracellular membrane trafficking (Hutagalung and Novick, 2011; Stenmark, 2009). Endocytic-membrane recycling is critical for both cell division and cytokinetic abscission. In cell culture, Rab11 positive recycling endosomes coordinate cell cycle progression and mitotic spindle positioning (Hehnly and Doxsey, 2014). Recycled membrane is trafficked to the midbody and impaired Rab11 or Rab35 endocytic-recycling pathways result in delayed or failed abscission (Dambournet et al., 2011; Frémont et al., 2018; Frémont et al., 2017; Kouranti et al., 2006). For example, disrupting Rab11 directed trafficking perturbs Kupffer’s vesicle morphogenesis during embryonic development in the zebrafish (Rathbun et al., 2020).

Here we characterize Rab25, an epithelial specific membrane recycling protein, that is a member of the Rab11 subfamily (Goldenring et al., 1993; Mitra et al., 2017). Rab25 has been implicated in cancer cell metastasis (Caswell et al., 2007; Mitra et al., 2017), but its role in embryonic tissue morphogenesis has not been examined. We demonstrate that maternal-zygotic Rab25a and Rab25b mutant embryos exhibit epithelial spreading delays. Here we propose an EVL endocytic-recycling defect in mutant embryos resulted in delayed or failed intercellular bridge abscission during cytokinesis which slowed epiboly movements. Failure of timely bridge scission resulted in the progressive formation of multinucleate cells which remained mitotically active, resulting in heterogenous cell sizes with anisotropic cell shapes and spatial arrangements in the EVL. Abnormal EVL cells exhibited an overall reduction in cortical actin density during epiboly. Sparse actomyosin networks were associated with uncoordinated cell behaviours, balanced tensions and altered viscoelastic responses during epiboly.

## Results

### Fluorescently tagged Rab25 localizes near the plasma membrane, centrosomes and cytokinetic midbody

In zebrafish, *rab25a* and *rab25b* transcripts are maternally deposited and their levels increase during gastrulation (https://www.ebi.ac.uk/gxa/home). Whole-mount *in situ* hybridization revealed ubiquitous expression of both genes in the blastoderm prior to epiboly initiation, followed by EVL restricted expression during epiboly (Figure 1A). EVL expression is consistent with reported epithelial specific function of Rab25 in mammals (Goldenring et al., 1993).

**Figure 1.**
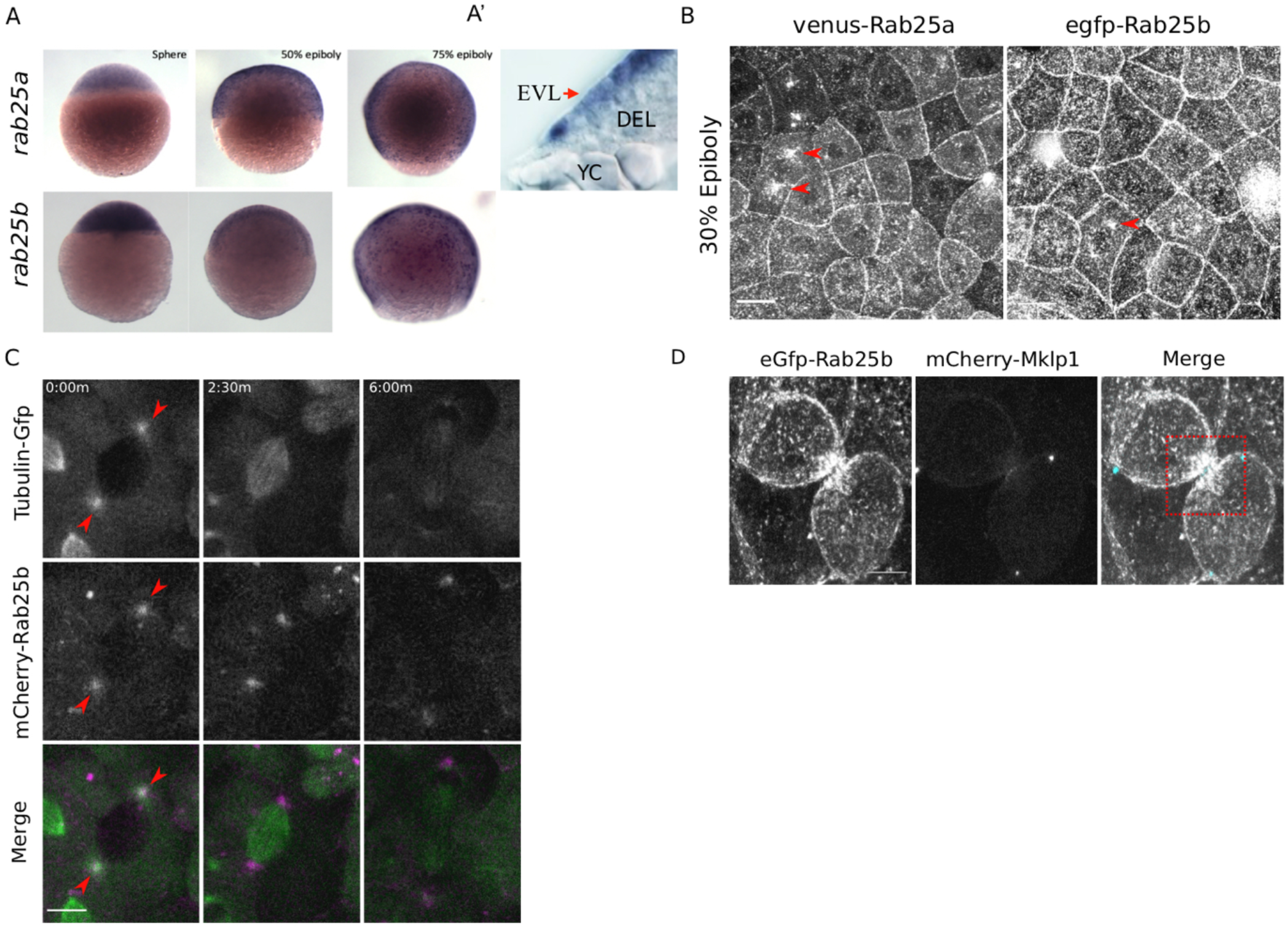
*rab25a* and *rab25b* expression pattern and subcellular localization. (A) Bright field images of whole-mount *in situ* hybridizations for *rab25a* [top row] and *rab25b* [bottom row], in WT embryos; lateral views with animal pole positioned to the top. (A’) Section of a WT embryo showing *rab25a* expression restricted to the EVL (arrow,top row far right panel); transcripts absent from the deep cells (DEL) and yolk cell (YC). (B) Confocal z-projections of venus-Rab25a and eGfp-Rab25b subcellular localization in WT embryos; red arrowheads denote perinuclear aggregates. Scale bar, 20μm. (C) Confocal z-projections of stills from time-lapse movies of transgenic Tubulin-GFP (green) embryos expressing mCherry-Rab25b mRNA (magenta). Arrowheads denote co-localization of mCherry-Rab25b at centrosomes. Scale bar, 20μm. (D) Confocal z-projections of eGfp-Rab25b (white) and mCherry-Mklp1 (teal) localization in WT embryos; box highlights enrichment of eGfp-Rab25b adjacent to the midbody. Scale bar, 20μm.

While Rab25 has not previously been implicated in cytokinesis, the closely related Rab11 has direct roles in cytokinetic bridge stability and abscission (Rathbun et al., 2020). Given the similarities between Rab25 and Rab11 GTPases (Mitra et al., 2017), we hypothesized Rab25 may have similar roles in the zebrafish EVL during epiboly. To examine the dynamic subcellular distribution of Rab25a and Rab25b in the EVL, N-terminally fluorescently tagged Rab25a and Rab25b constructs were expressed in WT embryos by RNA injection at the 1-cell stage. By live confocal microscopy, Venus-Rab25a and eGFP-Rab25b were observed at the plasma membrane and in motile cytoplasmic puncta (Figure 1B; Video S1). Venus-Rab25a and eGFP-Rab25b appeared to localize near centrosomes during mitosis and then moved towards the opposing poles of dividing cells (Figure 1B; arrowheads; Video S1). Injection of *mcherry-rab25b* mRNA into transgenic animals expressing Tubulin-GFP (Fei et al., 2019) confirmed that mCherry-Rab25b localized near centrosomes and dissipated following cell division (Figure 1C; arrows). Recycling endosomes (REs) become positioned near centrosomes during mitosis, suggesting that Rab25 associates with REs in the EVL, as in other systems (Casanova et al., 1999; Hehnly and Doxsey, 2014).

Prior to abscission, eGFP-Rab25b dynamically localized within and at the membrane of intercellular bridges. Co-expression of eGFP-Rab25b with the midbody marker mCherry-Mklp1, demonstrated that eGFP-Rab25b became highly enriched adjacent to the midbody, the sites of bridge scission (Figure 1D; box) (Rathbun et al., 2020). Overall, the spatial pattern of tagged-Rab25a and Rab25b was comparable to Rab GTPases associated with recycling endosomes in other systems (Hardin et al., 2017; Hehnly and Doxsey, 2014; Kouranti et al., 2006; Mavor et al., 2016), suggesting that Rab25 may be involved in vesicle recycling during epiboly. Furthermore, distribution adjacent to the midbody was suggestive of a role in cytokinesis (Rathbun et al., 2020).

### Rab25a and Rab25b are required for normal epiboly movements

To explore the functions of Rab25a and Rab25b, CRISPR/Cas9 gene editing was used to generate maternal-zygotic (MZ) mutant lines. Guide RNAs were designed to target exon 2 which encodes the GTPase domain, the functional domain of Rab proteins (Figure S1A) (Mitra et al., 2017). We characterized two *rab25a* mutant alleles from two founder fish, a 13-base pair (bp) deletion (2.3) and 29-bp insertion (4). Each allele contained a premature stop codon that disrupted the GTPase domain. An 18-bp deletion was generated in *rab25b* which produced an in-frame mutation and deleted a portion of the GTPase domain (Figure S1A). Quantitative PCR analysis of transcript levels at shield stage showed undetectable levels of *rab25a* transcripts and reduced *rab25b* transcripts in *MZrab25a* and *MZrab25b* embryos, respectively (Figure S1B). Nonsense mediated decay can result in genetic compensation of similar sequences hence we examined the levels of *rab25a, rab25b* and *rab11a* in mutant embryos (El-Brolosy and Stainier, 2017; Rossi et al., 2015). *MZrab25a* embryos had elevated levels of *rab25b* transcript while *MZrab25b* mutants did not have elevated levels of *rab25a* transcript. *rablla* transcript levels were similar to WT in both mutants. Thus, *MZrab25a* mutants appeared to exhibit genetic compensation in the form of transcriptional adaption of *rab25b,* whereas *MZrab25b* mutants did not.

To examine the mutant phenotypes, WT, *MZrab25a* and *MZrab25b* embryos were time-matched and examined live by light microscopy. *MZrab25a and MZrab25b* embryos exhibited epiboly delays that worsened over time, with the greatest delays at late epiboly stages (Figure 2A,6-9hpf;2B). The epiboly delay in *MZrab25b* embryos could be rescued by a transgenic construct expressing full length *rab25b* under the control of a B-actin promoter, indicating that the phenotype results from the loss of Rab25b (Figure S1C). The mutant phenotypes apparently did not result from a general developmental delay, as the zebrafish organizer formed at the same time as time-matched WT embryos (Figure 2A, 6hpf; white asterisk). During epiboly, deep cells are positioned behind the leading edge of the EVL and do not move past the EVL margin (reviewed in Bruce & Heisenberg, 2020). Examination of deep cell and EVL epiboly revealed they were equally delayed, suggesting the blastoderm delay could be the result of an EVL specific defect (Figure S1D). Overall, we found that *MZrab25b* phenotypes were more severe than either of the *MZrab25a* alleles. Despite the epiboly delay, most *MZrab25a* and *MZrab25b* embryos completed gastrulation and most embryos survived to adulthood.

**Figure 2.**
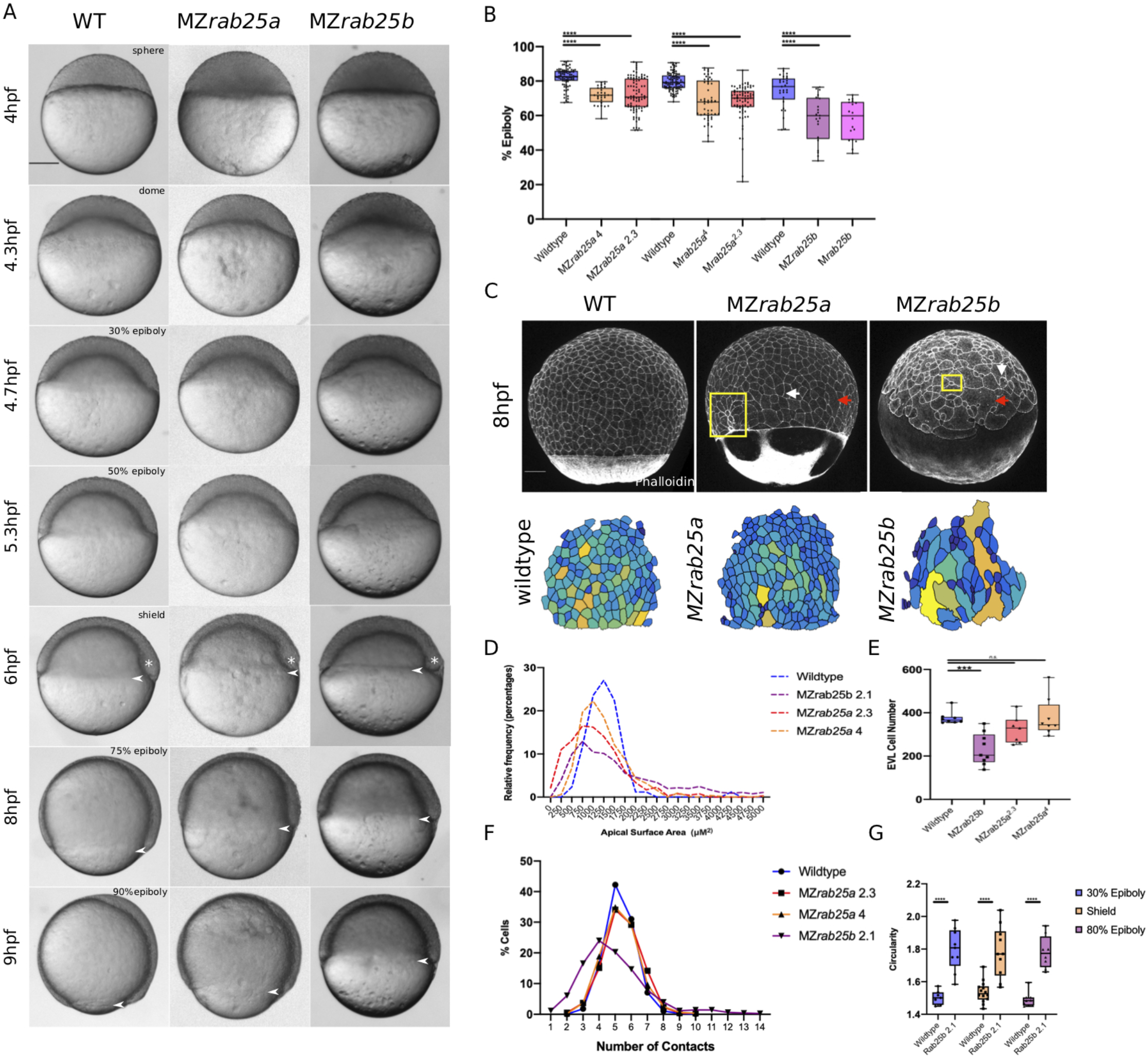
Epithelial spreading delays and heterogenous cell morphology, size and spatial arrangements in *MZrab25a* and *MZrab25b* embryos. (A) Time-matched bright field images of lateral views of WT, *MZrab25a* and *MZrab25b* embryos during epiboly. Arrows indicate blastoderm margin, asterisks denote embryonic organizer (shield). (B) Quantification of epiboly progression after 8hpf in: WT (n=80), *MZrab25a* (2.3, n=87) *MZrab25a* (4,n=28);WT (n=97), *Mrab25a* (2.3,n=73), *MZrab25a* (4,n=49); WT (n=29),MZ*rab25b* (n=21), *Mrab25b* (n=18). means: SEM;Two-Way ANOVA; ***p<0.001, ****p<0.0001.(N=3). (C) Confocal z-projections of time-matched lateral views of WT, *MZrab25a* and *MZrab25b* embryos at 8hpf stained with phalloidin and corresponding apical surface area heat maps. Cooler colours represent smaller areas, warmer colours represent larger areas. Yellow boxes indicate cells with reduced apices. Red arrows denote cells with increased apical surface area s. White arrows indicate curved cell junctions. Scale bar, 100 μm (D) Frequency distribution of apical surface areas of WT (n=817; N=8), *MZrab25a* 2.3 (n=651; N=14), *MZrab25a* 4 (n=654, N=15) and *MZrab25b* (n=503; N=15) embryos at 6hpf. (E) EVL Cell number in WT (n=8), *MZrab25b* (n=8), *MZrab25a* 2.3 (n=9) and *MZrab25a* 4 (n=7) embryos at 6hpf. Means:SEM;Two-Way ANOVA; ***p<0.001. (F) Frequency distributions of EVL cellular contacts number at 6hpf in (n=817; N=8), *MZrab25a* 2.3 (n=651; N=14), *MZrab25a* 4 (n=654, N=15) and *MZrab25b* (n=503; N=15). (G) Circularity quantifications for WT and *MZrab25b* embryos during epiboly. 30% epiboly: WT (n=7), *Mrab25b* (n=9); Shield: WT (n=14), *Mrab25b* (n=10): 80% epiboly; WT (n=9), *Mrab25b* (n=7). Mean;SEM; Two-Way ANOVA; ****p<0.0001.

We examined patterning and germ layer specification by in situ hybridization. EVL differentiation is required for normal epiboly (Fukazawa et al., 2010) and we found that expression of the EVL marker *keratin4 (krt4)* was similar in mutant and WT embryos (Figure S1E). Analysis of the mesoderm marker, *goosecoid,* and the endodermal marker *sox17,* also showed normal expression patterns (Figure S1D). Occasionally, *sox17* staining showed the dorsal forerunner cluster was disorganized in MZrab25a and MZrab25b embryos (Figure S1D; red arrows). Overall, germ layer specification appeared to be largely normally in mutant embryos. Together these data suggested that Rab25a and Rab25b may have a specific role in epiboly during zebrafish gastrulation.

### Cell shape and rearrangements indicative of epithelial defects in *MZrab25a* and *MZrab25b* embryos

The EVL restricted expression of *rab25a* and *rab25b* pointed to a primary defect in the EVL, consistent with the epithelial specific role of Rab25 in mammals (Goldenring et al., 1993; Jeong et al., 2019). EVL cell morphology was examined in stage-matched wild type and mutant embryos during epiboly and EVL cell shapes were largely normal at sphere stage. By shield stage in *MZrab25b* embryos, a range of EVL cell morphology were observed that were maintained into 75% epiboly (Figure S1H,H’). *MZrab25b* mutants exhibited more prominent EVL defects in the form of greater cell size heterogeneity and shape anisotropy compared to *MZrab25a* mutants (Figure 2C). The cell morphology defects in *MZrab25a* embryos were predominately confined to marginal regions (Figure 2C).

Our examination of phalloidin stained embryos, revealed that yolk cell actin networks were also perturbed in *MZrab25a* and *MZrab25b* embryos, suggesting yolk cell defects may contribute to the phenotypes (Figure 2C). However, depolymerizing actin in the yolk cell of wildtype embryos failed to cause similar EVL defects (Figure S1G). Although we cannot exclude the possibility of yolk cell actin defects contributing to the mutant epiboly delay, our data suggested the EVL morphology defects to be cell autonomous. Therefore, we focused our analysis on the EVL of *MZrab25a* and *MZrab25b* mutants.

Stage-matched embryos at 60% epiboly were used to quantify the EVL defects. This stage was chosen because that was how far epiboly progressed in most mutants by 8hpf and it was also when embryos exhibited the largest delays (Figure 2A,B). EVL mean surface area measurements, heat maps and frequency distributions were generated to assess the surface areas of cells in phalloidin stained mutant and WT embryos. *MZrab25a* and *MZrab25b* embryos exhibited heterogenous cell surface areas (Figure 2C,D). In WT, EVL cells had apical surface areas between 750-1750μm, while the distribution of EVL surface areas in *MZrab25a* and *MZrab25b* mutant embryos was broader, indicating both smaller and larger cell apices (Figure 2D).

Heterogeneity of apical surface area in mutants was associated with anisotropic cell shapes and disorganized spatial arrangements. While WT cells were pentagonal or hexagonal, as expected, *MZrab25a* and *MZrab25b* cells exhibited cell shape deviations and abnormal numbers of neighbours (Figure 2F,G). This was reflected by circularity measurements and cell-cell contact number, with larger circularity values indicating increased cell shape anisotropy (Figure 2G). Cells with increased apical surface area tended to have increased junctional curvature (Figure 2C; red arrows) whereas cells with reduced apices tended to be round (Figure 2C; yellow boxes). Notably, normally sized cells in mutants still displayed anisotropic cell shapes and nonlinear cell contacts (Figure 2C; white arrows). Compared to WT embryos, the EVL defects were associated with reduced cell number in *MZrab25b* but not *MZrab25a* embryos (Figure 2E). Embryo sizes were unchanged in *MZrab25a* and *MZrab25b* mutants (Figure S1F).

Cell size is reported to scale with nuclear number (Cao et al., 2017). Thus, we examined whether mutant cells with large apical surface areas were multinuclear by labeling WT, *MZrab25a* and *MZrab25b* embryos with membrane-GFP and the nuclear marker H2A-RFP. Large cells in *MZrab25a* and *MZrab25b* embryos were bi-or multinucleate (Figure 3A; red arrows), thus cell size scaled with nuclear number (Figure 3B). Overall, multinucleate large cells suggested cytokinesis failures in *MZrab25a* and *MZrab25b* embryos (Rathbun et al., 2020), consistent with the emergence of EVL defects following epiboly initiation, when the EVL is most proliferative (Campinho et al., 2013).

**Figure 3.**
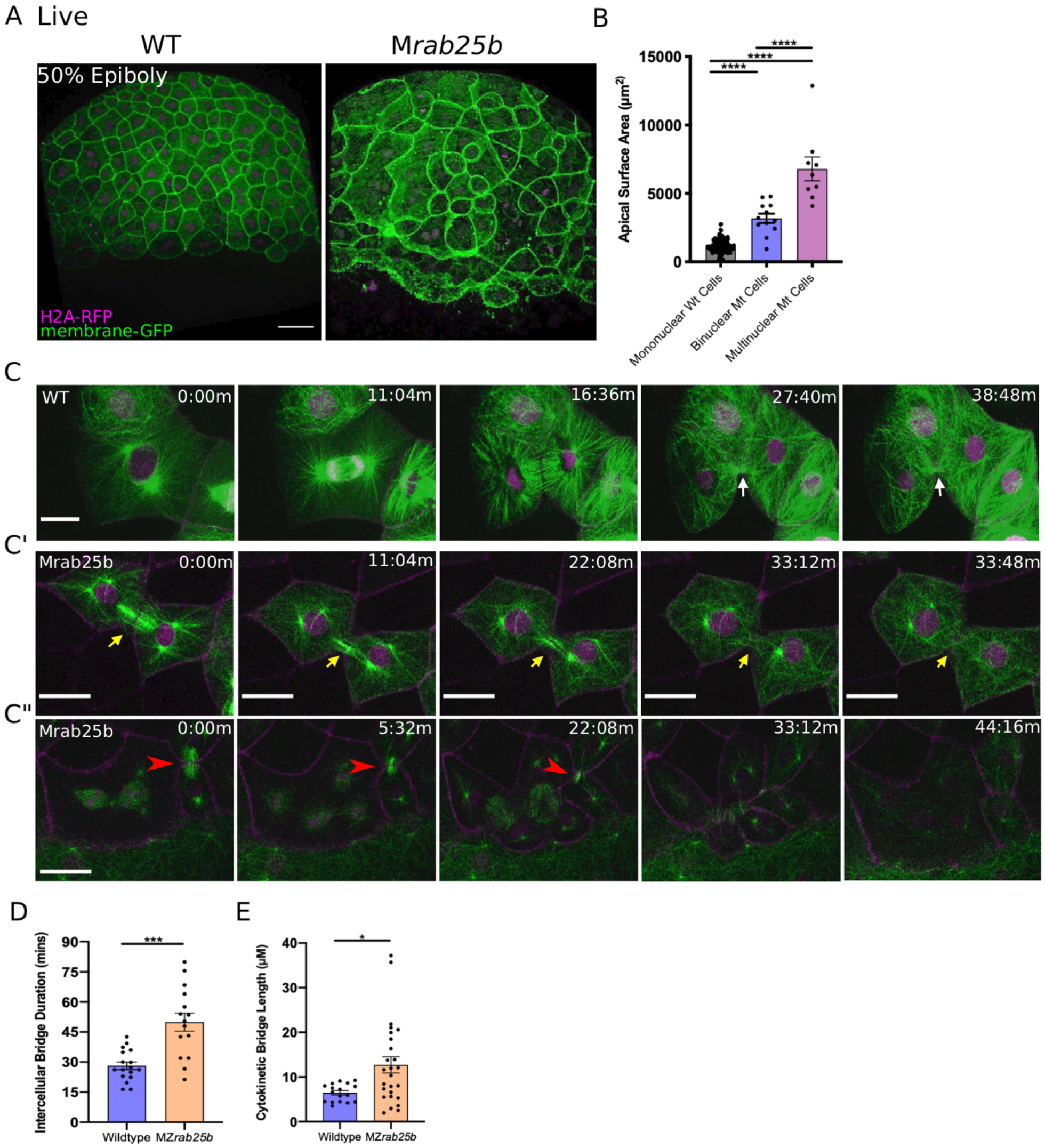
Unstable cytokinetic bridges and delayed abscission in *MZrab25b* embryos. (A) Confocal z-projections of stage-matched WT and *MZrab25b* embryos at 50% epiboly expressing mGFP (green) and H2A-RFP (magenta). Embryos positioned laterally, arrows denote bi- and multinucleate cells. Scale ba, 40μm. (B) Apical surface area quantifications of WT mononucleate (n= 151, N=6). *MZrab25a* and *MZrab25b* data was pooled: mutant bi-(n=12,N=6) and multinucleate cells (n=9,N=4) at 80% epiboly. Mean:SEM; One-Way ANOVA; ****, p<0.0001. (C-C”) z-projection of stills from confocal time-lapses of WT and *MZrab25b* embryos labelled for microtubules (green), nuclei (magenta) and plasma membrane (magenta) starting at dome stage; white arrow marks WT bridge; yellow arrows marks bridge regression in *MZrab25b* cell. (C”) Red arrowhead indicates cytokinetic bridge torn open by neighbouring morphogenic stress. Scale bar, 20μm. (D) Intercellular bridge duration in WT (n=17, N=3) and *MZrab25b* (n=15, N=4) embryos during epiboly initiation in EVL marginal regions. Mean:SEM; Mann-Whitney Test; ***, p<0.001. (E) Intercellular bridge length following formation in WT (n=17,N=3) and *MZrab25b* (n=26,N=4) embryos during epiboly initiation in EVL marginal regions. Mean:SEM; Mann-Whitney Test; *p<0.05.

### Cytokinetic Abscission defects during epiboly in *MZrab25b* mutants

Localization of eGFP-Rab25b near the cytokinetic midbody and the presence of large multinuclear EVL cells in *MZrab25a* and *MZrab25b* embryos indicated that Rab25 may function in cytokinesis. To investigate potential abscission defects in Rab25 mutants, we focused on *MZrab25b* embryos because they exhibited more severe defects than *MZrab25a* embryos. Cytokinetic bridges were characterized by labelling microtubules, nuclei and membrane in WT and mutant embryos. Following mitosis in WT cells, equatorial midzone microtubules were organized into apical cytokinetic bridges (Figure 3C; white arrows; Video S2). Initially, cytokinetic bridges underwent a fast phase of bridge narrowing and shortening, over approximately 10-12 minutes (Figure 3C; t=16:36m-27:40m). Intercellular bridges then remained connected for an additional 15-20 minutes, in which bridge length did not change significantly before abscission (Figure 3C; t=27:40-38:48min).

During mononucleate cytokinesis in *MZrab25b* embryos, the plasma membrane at the cleavage furrow underwent ingression similar to WT embryos (Figure 3C’, C”). Following cleavage, two types of abscission defects were detected. The first occurred during the initial, fast phase of bridge formation when bridge regression occurred in a small proportion of mutant cells during cytokinesis (5/23), leading to the formation of binucleate cells (Figure 3C’; yellow arrows; Video S3). In these instances, *MZrab25b* embryos expressing the midbody marker mCherry-Mklp1, showed a failure of mCherry-MKlp1 coalescence in cytokinetic bridges, which preceded bridge regression (Video S4; WT [left panel]; MZrab25b [right panel]).

The second defect occurred later during the abscission process. In WT embryos, daughter cells remained interconnected for ~26.23 mins before bridge scission (n=17), while in mutants bridges persisted almost twice as long (Figure 3D) (~50.35 mins; n=15). Bridges appeared to shorten and narrow normally during the initial phases of bridge formation in *MZrab25b* embryos and they were on average longer compared to wildtype (Figure 3C”; red arrowhead; 3E). While some bridges were still present at the end of the time-lapses, most long-lasting intercellular bridges appeared to be pulled open by morphogenetic stress in adjacent EVL regions, such as mitosis or basal cell extrusion, resulting in multinucleation (Figure 3C”; red arrowheads; Video S5). Overall, these observations are similar to *in vitro* work showing intracellular membrane trafficking regulates both cytokinetic bridge stability and timing of abscission (Dambournet et al., 2011).

### Multipolar Cytokinesis in *MZrab25a* and *MZrab25b* embryos

Binucleate cells remained mitotically active in *MZrab25b* embryos, resulting in continuous cytokinesis failures which produced multinucleated cells. Notably, in *MZrab25b* embryos, spindles in mono- and multinucleated EVL cells deviated from both the long axis of the cell and the animal-vegetal axis (Figure S2A). Thus, misoriented spindles likely contributed to the disorganized EVL cell arrangements (Campinho et al., 2013). During multipolar cytokinesis, both cleavage and abscission failures were seen. In some multinucleated cells, the plasma membrane appeared to snap back following furrow ingression, suggesting that the f-actin cytokinetic ring might be unstable (6/21) (Figure S2D). To investigate actin dynamics during cytokinesis, embryos were co-injected with RNAs encoding membrane-RFP and GFP-Utrophin. Cytokinetic rings progressed from basal to apical, as in other vertebrate epithelia (Figure 4A,B) (Higashi et al., 2016). During multinucleate cell cleavage in mutant embryos, we often observed failed contraction of the actin cytokinetic ring of EVL cells followed by cytokinesis failures (Figure 4C).

**Figure 4.**
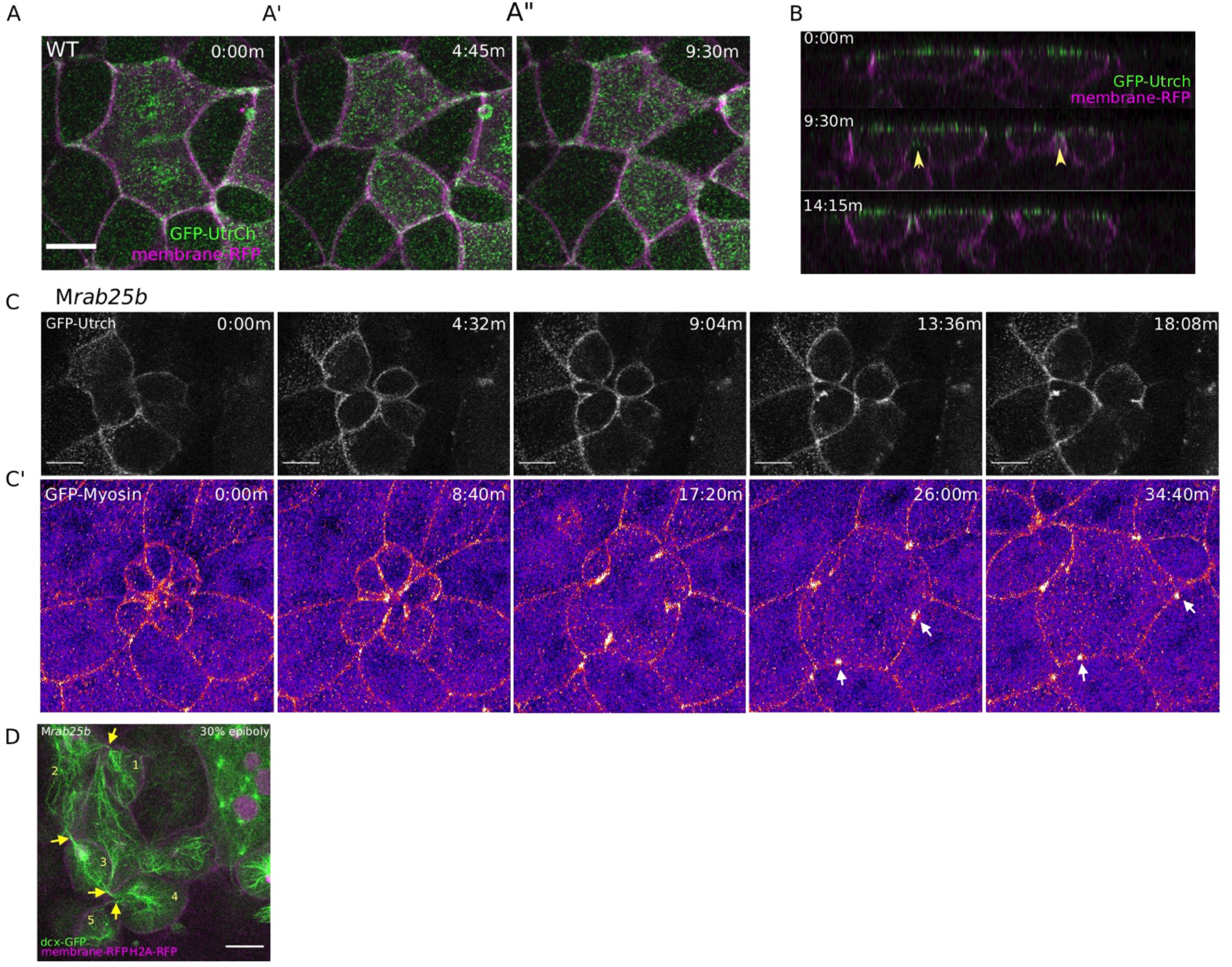
Multipolar Cytokinesis Failure in *MZrab25b* embryos. (A-A”) Confocal z-projections of stills from time-lapse of a WT EVL cell during mitosis injected with Gfp-Utrophin (green) and mRFP (magenta). Scale bar, 20μm. (B) Lateral views with apical to the top of stills from single-plane confocal time-lapses of WT EVL cells during mitosis expressing Gfp-Utrophin (green) and mRFP magenta). Arrowheads denote cleavage furrow ingression from basal to apical. (C) Confocal z-projections of stills from time-lapse of *MZrab25b* multipolar cleavage failure. F-actin labeled with Gfp-Utrophin. Scale bar, 20μm. (C’) Confocal z-projections of time-lapse of *MZrab25b* Tg (Myl1.1-gfp) (Fire-LUT) during multipolar cytokinesis failure. White arrows indicate myosin-gfp foci. (D) Confocal z-projection of *MZrab25b* embryo showing an array of EVL cells interconnected via cytokinetic bridges at 30% epiboly. Microtubules (green), nuclei (magenta) and plasma membrane (magenta), arrows and numbers denote connected cells and cytokinetic bridges. Scale bar, 20μm.

When multinucleate cells in *MZrab25b* mutants successfully completed cleavage (15/21), multiple daughter cells were generated that typically had reduced apical surface areas compared to surrounding cells (Figure S2D). Thus, successful cleavage during multipolar cytokinesis likely explains the presence of cells with reduced apical surface areas in mutant embryos. Most, but not all, daughter cells contained nuclei (not shown). Multipolar cytokinesis resulted in arrays of daughter cells interconnected through cytokinetic bridges across the EVL in *MZrab25b* embryos (Figure 4D; numerals and arrowheads; Video S6). As seen in mononuclear cell divisions, in multipolar divisions, intercellular bridges were not observed to undergo abscission but instead appeared to be torn open due to morphogenic stress, causing cytokinesis failures (Video S6). Following multipolar division failures, myosin-gfp foci were often observed along individual cell junctions, suggesting myosin was unevenly distributed along these EVL cell contacts (Figure 4C’; white arrows). Myosin-Gfp foci were apparently remnants of *de novo* tricellular vertices that are normally established along newly formed daughter cell interfaces, as observed in Xenopus during ectoderm cell divisions (Higashi et al., 2016).

We observed that cytokinetic intercellular bridge abscission failures resulted in the progressive formation of bi-and multinucleate cells during epiboly in *MZrab25b* embryos. The abscission defects disrupted EVL organization in mutant embryos, as cells exhibited variable sizes, shapes and number. Contributing to the tissue defects were basal cell extrusion events, cell-cell fusions and misoriented mitotic spindles (Figure S2E,F). Cytokinesis defects causing epithelial spreading delays aligns with previous work implicating cell division in epithelial cell rearrangements and epiboly (Campinho et al., 2013; Higashi et al., 2016).

### Marginal EVL cell rearrangements are disrupted during epiboly progression in *MZrab25b* embryos

The cellular defects in *MZrab25a* and *MZrab25b* embryos appeared largely to be the product of cytokinesis failures. We next sought to characterize cell behaviours during later epiboly stages, when the epiboly delay was greatest. During epiboly progression, marginal EVL cells elongate along the animal-vegetal axis and a subset of cells rearrange and intercalate into submarginal regions which enables the marginal circumference to narrow to eventually close the blastopore (Keller and Trinkaus, 1987; Köppen et al., 2016). To analyze EVL cell shape changes, live imaging of wildtype and mutant embryos expressing mRFP and Gfp-Utrophin was performed starting at 7 hpf.

In wildtype embryos, marginal EVL cells elongated as expected and a few cells changed shape by shortening their contacts with the yolk cell along the EVL margin (Figure 5A; green cells). Following these shape changes, 3-4 cells share one vertex with the underlying yolk cell (Figure 5A; red circle; t=25:26m), akin to a multicellular rosette. Resolution of rosettes in other contexts is linked to cell intercalation rates and tissue-shape changes (Blankenship et al., 2006; Zallen and Blankenship, 2008). Multicellular-yolk vertices resolved on average 22 minutes following their formation (Figure 5E). Resolution of marginal EVL rosettes resulted in *de novo* junctions formed between cells adjacent to intercalating cells as they exited into submarginal EVL regions (Figure 5A; orange cells; dotted line). Enrichment of cortical actin was observed when EVL cells intercalated into submarginal zones (Figure 5A’; Video S7 [left panel]).

**Figure 5.**
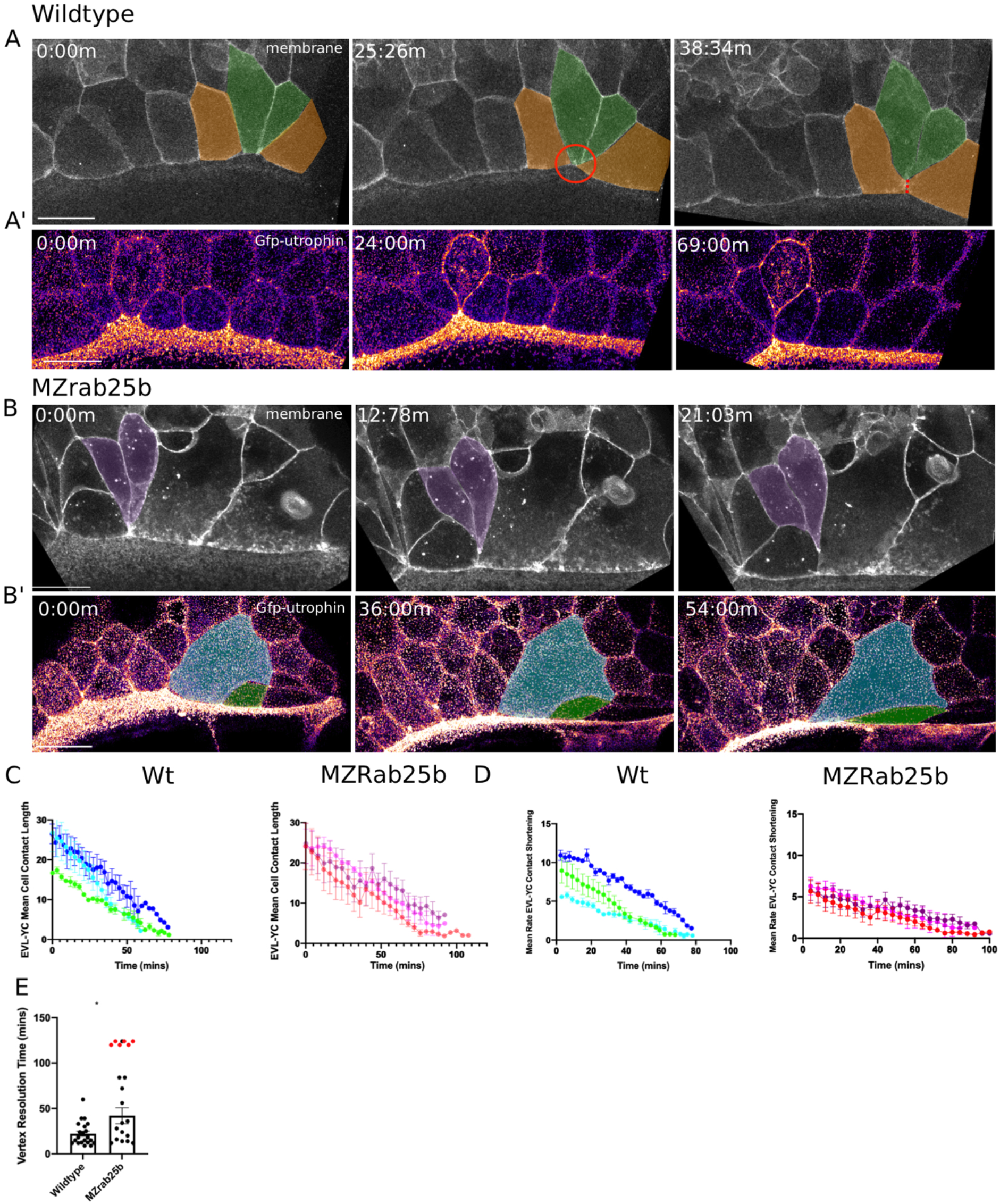
Aberrant EVL cell rearrangements in *MZrab25b* embryos. (A) Confocal z-projections of stills from a time-lapse of a WT embryo expressing mGFP starting at 7hpf. Lateral view focused on the margin. Green shaded cells shrink the EVL-YC junction and intercalate into submarginal zones. Orange shaded cells establish new cell-cell contacts following intercalation events (denoted by red dotted line). Circle denotes shared vertex with underlying yolk cell. Scale bar: 20μm (A’) Confocal z-projection time-lapse of WT embryo labelled for F-actin; (Fire-LUT) (Utrophin-Gfp) starting at 7hpf;lateral view; scale bar 20μm. (B) Confocal z-projection of stills from a confocal time-lapse starting at 7hpf of an *MZrab25b* embryo expressing membrane*-*Gfp. Purple shaded cells exit EVL marginal regions. Scale bar: 20μm (B’) Confocal z-projection of stills from time-lapse starting at 7hpf of an *MZrab25b* embryo labelled for F-actin (Fire-LUT) (Utrophin*-*Gfp); scale bar 20μm. Shaded cells denote an EVL circumferential stretching event. Scale bar: 20μm (C-D) EVL-YC mean contact length or shortening rate over time in rearranging EVL marginal cells in WT (N=3) and *MZrab25b* embryos (N=3). Mean:SEM. Each colour indicates a separate trial of a single embryo. Each line represents the average of the contact length or junction shrink rate in each trial (n=2-5). (E) Resolution times following formation of EVL-YC multicellular vertices. Mean:SEM. WT (n=20,N=4) and *MZrab25b* (n=12,N=5), unresolved *MZrab25b* vertices [red] (n=6,N=5). Mann-Whitney, *p<0.05.

Overall, EVL-YC junction shortening took longer in *MZrab25b* embryos compared to wildtype (Figure 5C). Additionally, while WT EVL-YC contact shrink rates were initially fast and slowed over time, *MZrab25b* contacts displayed more uniform, slower rates of shortening during epiboly (Figure 5D). Cells of all sizes were observed to shorten their yolk cell contacts in mutant embryos (Video S7 [right panel]). However, compared to wildtype, mutant cells with normal and large apical surface areas were slower to shrink their yolk cell contact, while cells with reduced apices did so more quickly (Figure S3A). This data suggests that the cell size defects impacted the rate of EVL cell shape changes, contributing to the slowed rate of epiboly in mutant embryos.

Resolution of multicellular-yolk vertices was significantly slower in mutant embryos compared to wildtype (Figure 5E). A large proportion of rosettes in marginal regions persisted for up to two hours following their formation (Figure 5E; red data points), preventing timely intercalation of cells into submarginal zones and potentially slowing overall EVL marginal circumference narrowing. Additionally, marginal rosettes in *MZrab25b* embryos appeared to deform the EVL-YC boundary, causing the margin to buckle animally, resulting in an uneven EVL margin (Figure S3B) and potentially hindering epiboly progression. Given normal sized cells exhibit both slow vertex resolution and intercalations rates compared to wildtype and that tagged-Rab25 constructs localized to vertices during marginal EVL rearrangement events (Video S8) suggests Rab25 may have an additional, direct role in promoting EVL cell intercalation events during epiboly.

Despite the epiboly delay in *MZrab25b* embryos, EVL cells were able to rearrange into submarginal zones in (Figure 5B; purple cells), with adjacent cells with large apices being stretched towards the site of intercalation (Figure 5B’; coloured cells). Marginal EVL cell stretching was in contrast to wildtype cell shape changes and would be predicted to slow the rate of epiboly, as cells normally maintain their shape or narrow and elongate along the embryonic AV axis (Köppen et al., 2006).

### Actomyosin network organization in MZ*rab25b* embryos

Our observations of EVL cells in mutants exhibiting misoriented and uncoordinated cell shape changes prompted us to examine f-actin and myosin in more detail given their fundamental roles in cell shape and rearrangements (Jewett et al., 2017). A gradual reduction in cortical actin over the course of epiboly was observed in phalloidin stained mutant embryos, which was first detected at 30% epiboly in *MZrab25b* embryos (Figure 6A-A”; Figure S4A). While cells of all sizes exhibited reduced F-actin signal intensity (Figure 6A; red arrows; Figure S6B), as cell size increased, cortical actin further decreased (Figure 6A; yellow arrows).

**Figure 6.**
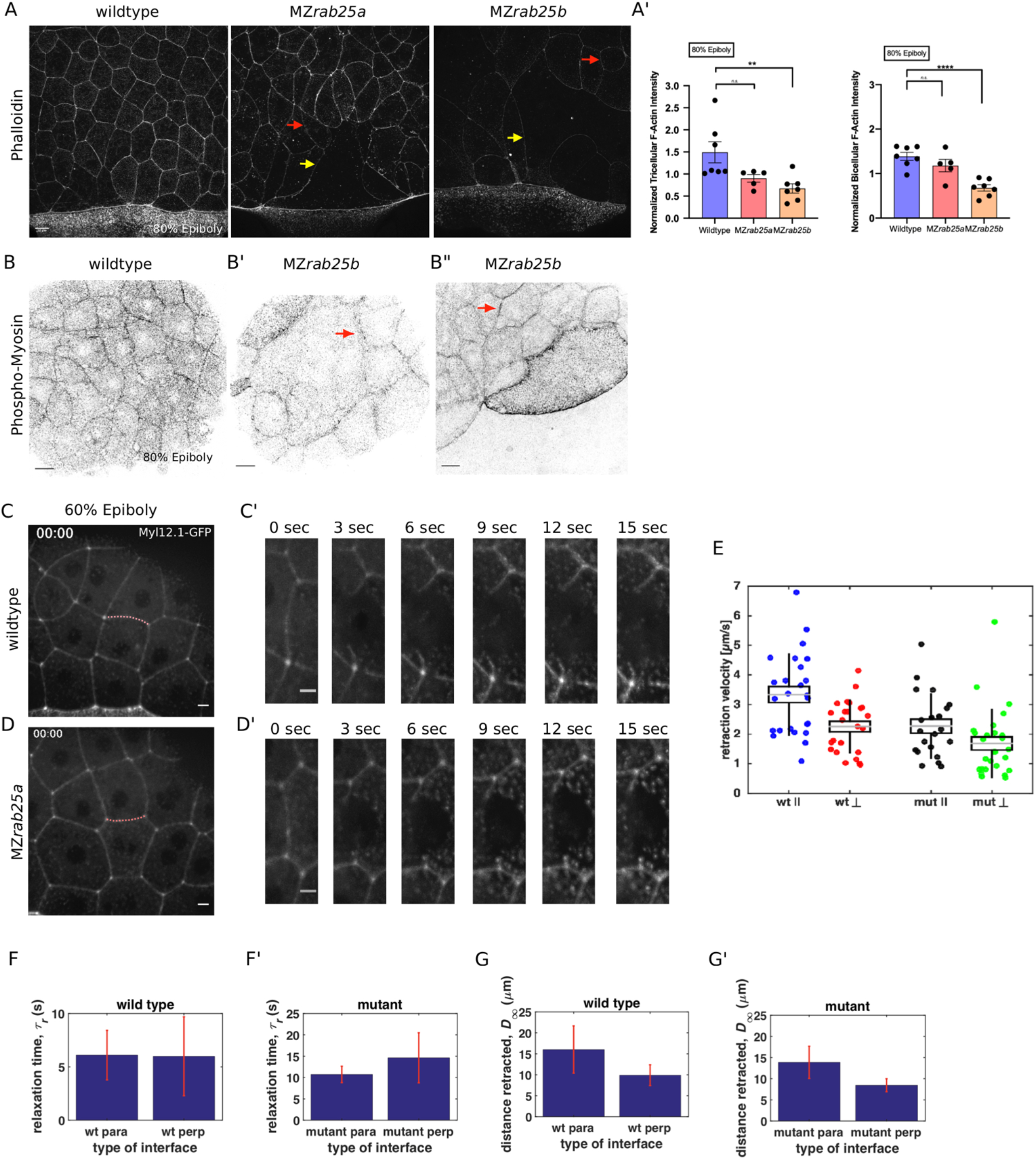
Reduced cortical actin and P-myosin associated with less tension and more viscous tissue responses in mutant embryos. (A) Lateral views with animal pole to the top of z-confocal projections of phalloidin stained WT, *MZrab25a* and *MZrab25b* embryos stage-matched at 80% epiboly; red arrows show reduced cortical actin in normal sized cells; yellow arrows show reduced actin in large cells; Scale bar 20μm. (A’) Quantification of normalized tricellular and bicellular f-actin intensity at 80% epiboly. WT (n=90,N=9), *MZrab25a* (n=90,N=9) and *MZrab25b* (n=90,N=9).; means;SEM; Mann-Whitney, **,p<0.001. (B) z-confocal projections of WT and *MZrab25b* embryos at 80% epiboly antibody stained for phospho-myosin. Red arrows denote uneven distribution of P-myosin along individual *MZrab25b* cellular junctions. Scale bar 20μm. (C-D’) z-confocal projections of WT or *MZrab25a* Tg(Myl1.1-Gfp) at 60% Epiboly; lateral positioned embryo focused on EVL margin; Red line marks the ablated junction. Scale bar 5μm. (E-G’) Initial recoil velocity, relaxation time and distance retracted in WT and MZrab25a embryos following junction laser cutting (see methods). WT and *MZrab25a* perpendicular and parallel cuts (n=26,23);(n=21,25).

Antibody staining for phosphorylated myosin (pMyosin), the active form of myosin and a proxy for contractility, revealed that most EVL cells in *MZrab25b* embryos exhibited reduced junctional pMyosin compared to wildtype embryos at 80% epiboly (Figure 6B,B’). In *MZrab25b* embryos, we occasionally observed some EVL cells with elevated levels of cortical pMyosin (Figure 6B”). Furthermore, while junctional pMyosin was uniformly distributed along individual cell-cell contacts in wildtype embryos (Figure 6A), in MZr*ab25b* mutants, it was often diffuse and fragmented along cell-cell contacts (Figure 6B’-B”; arrows). Overall, reduced cortical actin was associated with weak junctional pMyosin in *MZrab25b* embryos, with some localized increases in pMyosin in individual cells.

We next characterized the distribution of actin and myosin in live embryos. In *MZrab25b* embryos transgenic for Myosin light chain 12, genome duplicate 1-gfp (Myl12.1-GFP) myosin intensity was often weak at shield stage compared to wildtype and became heterogeneously distributed in the EVL, as some cells contained elevated levels of cortical myosin following aberrant shape changes (Video S9). GFP-Utrophin was also disorganized in *MZrab25b* EVL cells following aberrant cell behaviours (Video S6 [right panel]). Collectively, these data indicated that ectopic activation of actomyosin in localized EVL regions are likely the result of disorderly cell movements and rearrangements. This is similar to the accumulation of actomyosin in the Drosophila germband during unorganized cell rearrangements (West et al., 2017). We propose the abnormal cell shape changes and behaviours in MZrab25b mutants are likely the product of the overall reduction of contractile actomyosin networks and disrupted tissue architecture.

### Altered viscoelastic responses in *MZrab25a* mutant EVL cells

The initial recoil velocity of laser ablated junctions is a measure of tension (Hutson et al., 2003). Reduced cortical actin and pMyosin suggested decreased contractility in the EVL of *MZrab25a* and *MZrab25b* embryos. To examine this, junctions perpendicular and parallel to the EVL-YSL margin were ablated in WT and *MZrab25a* mutant embryos that were transgenic for Myosin-GFP. We analyzed cells near the EVL margin in embryos at 60-70% epiboly, when EVL tension is highest and epiboly was the most delayed in *MZrab25a* and *MZrab25b* embryos (Campinho et al., 2013). To eliminate potential effects of contact length on tension, only cell junctions with similar lengths were quantified.

In WT, cell vertices at the ends of contacts parallel to the margin recoiled faster than perpendicular vertices (Figure 6C,C’,E). Cells behave as viscoelastic materials; thus, tissue viscosity can impact junctional recoil velocities. Laser ablation results can be modelled as the damped recoil of an elastic fibre, using a Kelvin-Voigt element to represent the junction (Kumar, Ingber, 2006). In this context, the maximum distance retracted after ablation is proportional to the stress sustained by the junction, while the rate at which that maximum distance is attained (measured by a relaxation time) is proportional to the viscoelastic properties of the tissue. The relaxation times of parallel and perpendicular junctions following ablation were not significantly different, suggesting that the difference in initial recoil velocities can be attributed to differences in tension, not viscosity (Figure 6F). Stress modelling was consistent with the initial recoil velocities (Figure 6G). Elongating junctions perpendicular to the yolk cell having slow recoil velocity is similar to tensions reported for fluidizing, growing junctions during wound healing (Tetley et al., 2019). Overall, our data suggests that as the yolk-cell begins pulling the EVL between 60-75% epiboly, mechanical stresses are anisotropic, being highest along cell contacts oriented parallel to the EVL-YC boundary in marginal EVL regions.

In *MZrab25a* embryos, the initial recoil velocities of parallel and perpendicular junctions were significantly different, but this difference was diminished compared to wildtype embryos (Figure 6D-E). Additionally, both parallel and perpendicular junctions in *MZrab25a* mutants exhibited slower recoil velocities compared to wildtype (Figure 6E). Stress modelling was consistent with both these results (Figure 6G’). However, the relaxation time of the EVL following the initial recoil was much longer in mutant embryos compared to wildtype, suggesting a more viscous response following laser ablation (Figure 6F’).

Overall, this suggests that forces are more balanced in the EVL and that viscoelastic responses are defective in Rab25 mutant embryos. Anisotropic tensions and contraction drive epithelial wound closure in Drosophila and epithelial monolayers in culture (Zulueta-Coarasa and Fernandez-Gonzalez, 2018). Thus, balanced forces would be predicted to disrupt force patterns in the EVL that drive oriented cell shape changes required for tissue elongation during epiboly progression. The overall reduction in actomyosin density and contractile networks likely contributes to the relaxation of tensions within the EVL of mutant embryos. Importantly, force is propagated across tissues through the actomyosin cytoskeleton. Accordingly, reduced actomyosin and tension would also be predicted to impact force transmission across the EVL during epiboly, disrupting the rate of tissue spreading. Lastly, inhibition of cell divisions in amniotes has been shown to increase viscosity, lending support to our data suggesting an association between cytokinesis failures and increased tissue viscosity (Saadaoui et al, 2020). Increased viscosity would slow the rate of EVL deformation in response to yolk cell pulling forces, given viscous responses are slower than elastic deformations (Petridou and Heisenberg 2019b).

### *MZrab25a* and *MZrab25b* mutants exhibit endocytic trafficking defects

Given the subcellular distribution of tagged-Rab25 and the reported role of Rab25 in cargo recycling in cell culture, we postulated that a vesicular trafficking defect may underlie the mutant phenotypes (Cassanova et al., 1999; Mitra et al., 2017). Recycling endosomes (REs) are membrane compartments that act as hubs for biosynthetic and endocytic material (van Ijzendoorn, 2006). Recycled membrane is trafficked from REs to the cell surface and towards the cytokinetic midbody. If Rab25 directs recycled membrane, Rab25 should co-localize with other members of the recycling pathway like Rab11, a conserved recycling endosome protein in multiple systems (Goldenring, 2015). In addition, an accumulation of material trapped in RE compartments would be expected in mutant embryos, which could result in increased size and/or number of RE compartments (Mavor et al.,2016; Langevin et al., 2005).

To examine Rab25’s role in recycling, Venus-Rab25a or eGFP*-*Rab25b were co-expressed with mCherry*-*Rab11a and found to partially co-localize during epiboly, suggesting that Rab25 localizes to REs (Figure7A; S5A). To examine RE number and size, antibody staining for Rab11b was performed on WT and mutant embryos. In WT embryos, Rab11b staining was junctional and cytoplasmic, similar to mCherry-Rab11a (Figure 7B). Co-staining for Rab11 and Rab25 was not possible, due to the lack of a zebrafish Rab25 antibody. In *MZrab25b* embryos an accumulation of large Rab11b positive endosomes was observed that were not present in WT embryos (Figure 7B). Notably, in mutant embryos these endosomes appeared in cells with both large and small apices, suggesting that the trafficking defects occurred irrespective of cell size. We next wanted to determine the origin of the material trapped in Rab11b endosomes.

**Figure 7.**
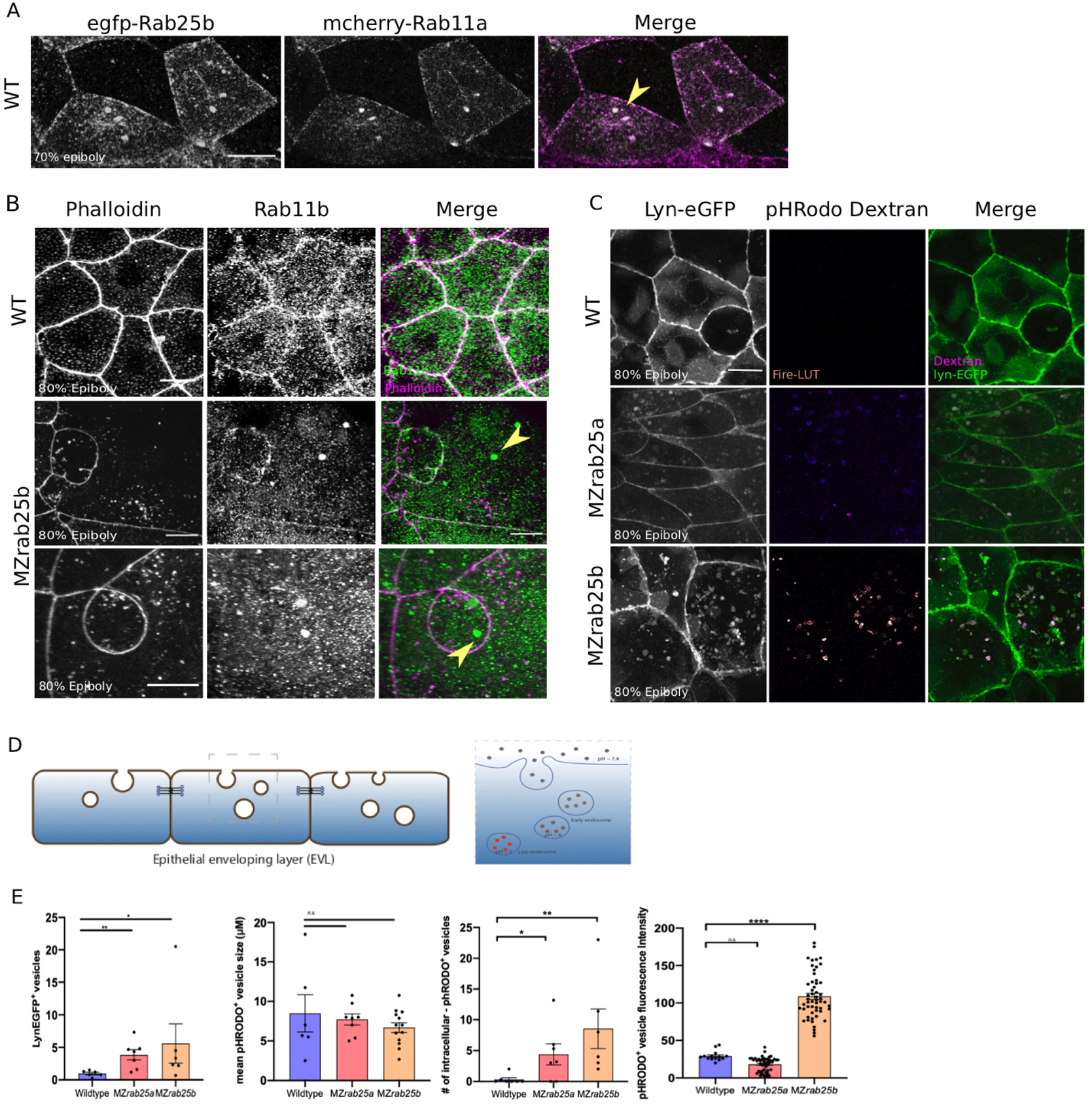
Tagged-Rab25 and mcherry-Rablla co-localize and *MZrab25a/b* exhibited RE trafficking defects. (A) z-projection stills of a wildtype embryo at 80% epiboly co-expressing mCherry-Rab11a (green) and eGfp-Rab25b (magenta); arrowheads denote overlap. (B) Rhodamine phalloidin stained (magenta) and Rab11b antibody (green) stained WT and *MZrab25b* embryos at 80% epiboly; arrowheads denote large Rab11b endosomes. Scale bar; 20μm (C) Live WT, *MZrab25a* and *MZrab25b* embryos expressing lyn-EGFP (green) and containing cytoplasmic phRODO dextran (magenta) following incubation. Scale bar: 20μm. (D) Schematic of phRODO dextran apical endocytosis (E) Mean # of lynEGFP- or pHRODO-positive vesicles/cell. WT, *MZrab25a* and *MZrab25b* embryos (N=7,7,6). Fluorescence intensity measured over a 1μm line in phRODO-positive vesicles; WT (n=15); *MZrab25a* (n=46); *MZrab25b* (n=54). Mean surface area of phRODO vesicles; WT(n=6); *MZrab25a and MZrab25b* (n=23,13). Means; SEM; significance using Mann-Whitney test. Scale bars;(A-C) 20μm

While recycling endosomes contain both newly synthesized and endocytosed cargo, Rab25 is thought to primarily traffic recycled membrane (Mitra et al., 2017). Therefore, we hypothesized that REs in mutants would contain trapped endocytic material. To analyze this, Lyn-eGFP expressing WT and *MZrab25a/b* embryos were incubated in pHRODO dextran. The fluorescently tagged peptide Lyn eGfp labels intracellular membrane compartments during trafficking (Sonal et al., 2014). phRODO dextran can only be internalized by EVL cells through apical endocytosis (Figure 7D). phRODO fluorescence increases in acidic environments, indicating endo-lysosomal compartments.

In WT cells, we detected a small number of Lyn-eGFP cytoplasmic vesicular bodies with faint phRODO dextran signal (Figure 7C). In contrast, both Lyn-eGFP and pHRODO positive vesicles accumulated in *MZrab25a/b* embryos, consistent with defects in apical-endocytic membrane recycling (Figure 7C,E). We also observed reduced junctional Lyn-eGFP in mutants compared to wildtype, perhaps representing reduced fusion of vesicles with the plasma membrane. Notably, the greatest accumulation of intracellular vesicles was observed in cells with large apices. Vesicles also accumulated in cells with normal morphologies and small apices, although to a lesser extent. These findings suggested that the trafficking defects at least partially reflected the loss Rab25 cargo recycling in mutant embryos as opposed to a secondary effect from changes in EVL cell morphology. A proportion of the intracellular vesicles were acidic, as shown by increased fluorescence intensity of vesicles in in *MZrab25b* embryos (Figure 7E), perhaps representing late endosomes or lysosomes (Langemeyer et al., 2018) and reflecting the downstream consequences of an overall trafficking disruption. These finding suggest that disrupted recycling likely explains the EVL morphogenesis defects in *MZrab25* mutants.

## Discussion

Rab GTPases comprise the largest family of small GTPases and are an important class of membrane trafficking regulators (reviewed in Nassari et al, 2020). Recently, several Rabs have been shown to play roles in morphogenesis and embryonic development (Jewett et al. 2017; Ossipova et al., 2014, 2015; Rathbun et al., 2020). Rab25 is reported to be epithelial specific in mammals and is a member of the Rab11 subfamily (Casanova et al., 1999; Goldenring et al., 1993), which has been implicated in apical recycling, cytokinesis, junction reinforcement and turnover (Iyer et al., 2019; Langevin et al., 2005; Rathbun et al., 2020; Woichansky et al., 2016). Rab25 mutant mice are viable but exhibit defects in skin homeostasis suggesting role in epidermal formation (Nam et al., 2010; Jeong et al., 2019). The function of Rab25 during development is largely unexplored and here we present the first characterization of two *rab25* genes in the zebrafish embryo.

Zebrafish *rab25a* and *rab25b* are expressed in the surface epithelial layer, the EVL, during gastrulation. *MZrab25a* and *MZrab25b* mutant embryos exhibit epiboly delays and EVL cellular morphology defects. As described further below, we propose that the heterogeneous EVL cell sizes in mutant embryos, resulting from cytokinesis failures, impairs epiboly in two ways; by hindering local cell rearrangements and altering the viscoelastic properties of cells.

### Role for Rab25 in Cytokinetic Abscission

The overall subcellular distribution of Rab25 constructs is consistent with its function as a recycling protein (Hehnly and Doxsey, 2014; van Ijzendoorn, 2006; Langevin et al. 2005). Rab25 constructs were highly motile in the cytoplasm as puncta and transited towards the plasma membrane, suggestive of recycling events. During mitosis, Rab25 localized near centrosomes, where recycling endosomes are positioned to direct membrane to the intercellular bridge during cytokinesis (Frémont and Echard, 2018). Lastly, Rab25 co-localized with the known recycling protein Rab11.

*MZrab25b* embryos exhibited a range of cytokinesis defects during epiboly which led to the formation of bi-and multinucleate cells. Unstable cytokinetic bridges were associated with failed formation of cytokinetic midbodies while persistent apical cytokinetic bridges experienced either failed or delayed abscission. Binucleate cells remained mitotically active generating large, multinucleate cells with reduced cortical actin which continued to exhibit abscission defects leading to an array of interconnected cells across the EVL. These observations point to a previously uncharacterized role for Rab25 in cell division.

Cell division and cytokinesis are multicellular processes (Herszterg et al., 2013), and morphogenic stress in the form of neighbouring cell divisions or basal cell extrusion events appeared to tear open persisting cytokinetic bridges in mutant embryos. In cell culture, when abscission is delayed through siRNA knockdown of Rab35, mitosis similarly tears open cytokinetic bridges (Frémont et al., 2017). Our observations of a few cells failing to initially form bridges and a large proportion cells experiencing delayed abscission in *MZrab25b* mutants is highly similar to the phenotypes of Rab11 and Rab35 knockdown experiments *in vitro* (Frémont et al., 2017; Kouranti et al., 2006; Mierzwa and Gerlich, 2014). Furthermore, optogenetic clustering of Rab11 during Kupffer’s vesicle morphogenesis in zebrafish resulted in abscission failure (Rathbun et al., 2020). Therefore, our data implicates Rab25 in having a direct role in cytokinesis.

Current models of membrane dynamics during intercellular bridge cleavage suggest coordination amongst many Rab recycling pathways (Frémont and Echard, 2018). Rab11 and Rab35 are thought to act in a redundant manner in regulating bridge scission *in vitro* (Frémont et al., 2017), although whether this is the case in animal models has yet to be investigated. Our evidence suggests *rab25a* and *rab25b* have similar functions during epiboly. We propose this as *rab25b* transcripts were upregulated in *MZrab25a* embryos which was associated with less severe EVL cytokinesis defects.

Rab11 vesicle trafficking has been shown to be important for cytokinesis during zebrafish KV development (Rathbun et al., 2020). As KV is an EVL derived organ (Oteiza et al., 2008), it is likely Rab11 has an abscission function during epiboly. Previous Rab11 dominant negative or morpholino approaches have not produced phenotypes similar to *MZrab25b* mutant embryos, but this can likely be attributed to maternally expressed Rab11 family members rescuing depletion of zygotic Rab11 in these experiments (Clark et al., 2011). Rab11 and Rab25 may be acting stepwise in vesicle delivery towards the cytokinetic midbody to regulate abscission or may regulate separate trafficking pathways altogether. A more attractive hypothesis is that Rab11 and Rab25 work as a complex to regulate abscission or have partially overlapping trafficking pathways, a proposal consistent with both constructs co-localization and parallel dynamics during epiboly. Coregulation of Rab proteins on trafficking pathways is observed in *in vitro* models (reviewed in Stenmark, 2008).

An open question is why Rab25a or Rab11 cannot compensate for loss of Rab25b in *MZrab25b* mutant embryos. A possible explanation may be that more Rab25b protein is maternally deposited compared to Rab11a or Rab25a, thus, disruption of Rab25b without transcriptional adaptation of *rab25a* or *rab11a* may not be able to rescue the abscission phenotypes. Given the lack of Rab25b antibody this is difficult to determine. Membrane trafficking becomes increasingly important in morphogenetically active tissues, as a higher degree of plasma membrane remodelling occurs (reviewed in Pinheiro and Bellaiche, 2018). This is exemplified in the related teleost fish Fundulus, where membrane turnover in the EVL was shown to steadily increase as epiboly progressed (Fink and Cooper, 1996), suggesting that membrane recycling may similarly increase. Thus, having high amounts of Rab trafficking molecules during epiboly is most likely needed for normal cytokinesis.

### Cytokinesis Failures disrupts EVL morphogenesis

Cell division appears to have two major roles in epithelial morphogenesis. Cell division mediated intercalations (CMI) power epithelial rearrangements in chick, quail and Xenopus embryos during gastrulation (Firmino et al., 2016; Higashi et al., 2016; Saadaoui et al., 2020). In general, cell rearrangements have well defined roles in tissue morphogenesis and embryonic development (reviewed in Zallen and Blankenship 2008). Thus, failed CMI disrupts rearrangements critical for tissue development, as shown in amniote gastrulation and mouse limb bud development (Firmino et al., 2016; Lau et al., 2015; Saadaoui et al., 2020; Wyngaarden et al., 2010). A second role of cell divisions is the regulation of tissue fluidity and viscoelasticity (reviewed in Petridou and Heisenberg 2019b). Recent experiments using laser ablation and micropipette assays in amniote epiblast cells and zebrafish deep cells have shown that inhibiting cell division increases tissue viscosity, effectively slowing tissue shape changes (Petridou et al. 2019a; Saadaoui et al. 2020). Our data in *MZrab25a* and *MZrab25b* embryos suggests that failed cytokinesis disrupted cell intercalation events during epithelial epiboly and increased tissue viscosity, with both likely contributing to slow EVL spreading.

Marginal epithelial cell rearrangements are required for normal epiboly movements of surface epithelia in zebrafish, Fundulus and Tribolium embryos (Jain et al., 2019; Keller and Trinkaus, 1987; Köppen et al., 2006). Similar to T1 transitions and rosette resolution described in the Drosophila germband (Blankenship et al., 2006), marginal EVL cell intercalations bring into contact initially non-neighboring cells. Marginal intercalation leads to EVL circumference narrowing needed to close the blastopore (Köppen et al., 2006). The EVL cell shape heterogeneity in mutant embryos led to an overall slowing of marginal cell intercalation events, which likely impaired EVL circumference narrowing and the rate of epiboly in *MZrab25a* and *MZrab25b* embryos. While we investigated cell rearrangement events predominantly in EVL marginal regions, our time-lapses of wildtype embryos revealed EVL cell neighbour exchanges throughout most regions of the epithelium, which to date has not been reported.

### Rab proteins and cytoskeletal dynamics

Tension anisotropy promotes tissue shape changes during morphogenesis. For example, heterogenous contractile networks are required for timely wound healing and apical cell extrusion (Le et al., 2020; Zulueta-Coarasa and Fernandez-Gonzalez, 2018). Our laser cutting analysis revealed polarized tensions within the EVL, with cell junctions aligned parallel to the EVL-YC boundary having high recoil velocities compared to perpendicular junctions elongating during epiboly progression. Higher tensions along parallel contacts indicate cells may dynamically narrow along the circumferential embryonic axis during late-phase epiboly, consistent with the tissue narrowing and elongating. It may also mean some junctions are resisting external forces to maintain their initial length. In support of our notion that tension heterogeneity drives EVL morphogenesis, the overall relaxation of tensions in mutant embryos disrupted polarized force patterns which was associated with disorganized and stochastic cell behaviours that hindered collective cell movements and epiboly.

Similar to what has been described in several different Drosophila epithelia (Lecuit et al., 2011), we found that abnormal cell shape changes were associated with a global reduction of cortical actomyosin networks in *MZrab25a* and *MZrab25b* embryos. We observed that enlarged, multinucleated cells had the largest depletions of cortical actomyosin. This reduction in cortical actomyosin provides a likely explanation for the significant slowing of cell rearrangements exhibited by multinucleate cells compared to neighbouring cells with normal or reduced apical surface areas. Furthermore, this finding aligns with the observation that greater epiboly delays were associated with the increased presence of multinucleated cells. Reduced cortical actin also explains the tension defects shown via laser ablation, which we also propose results in perturbed force transmission across the EVL. Despite the relationship observed between cell size and cortical actin density, cytokinesis failures do not explain weak actomyosin intensity throughout the EVL in normal sized cells in both mutant backgrounds. This suggests that Rab25 may regulate the cortical actin network independently of its role in abscission.

Rab proteins have conserved roles in actomyosin network maintenance. During Drosophila mesectoderm invagination and germband extension, Rab35 localizes to actomyosin networks and is required for sustained cell-cell interface contraction (Jewett et al., 2017). Disruption of Rab35 impairs cell shape changes and overall tissue morphogenesis in Drosophila (Jewett et al., 2017). In Xenopus bottle cell ingression and neuroepithelial folding, Rab11 similarly promotes actomyosin contractility to drive both developmental programs (Ossipova et al., 2014, 2015). Thus, Rab25 having a role in actomyosin regulation during epiboly is consistent with Rab proteins in other systems.

The mechanism for Rab25 cortical actomyosin maintenance is currently unclear and requires further study. Rab proteins promote cytoskeletal changes at subcellular sites by delivering actin remodelling proteins. For example, during abscission, Rab11 delivers p50RhoGAP to depolymerize intercellular bridge actin networks (Schiel et al., 2013). Additionally, a Rab11 effector, Nuf, is involved in actin polymerization during Drosophila cellularization (Cao et al., 2008). Rab25 vesicles may also contain lipids involved in signalling pathways which reorganize actin networks. For example, Rab35 vesicles contain the PtdIns(4,5)P2 phosphatase OCRL which remodels F-actin during abscission (Dambournet et al., 2011).

Alternatively, junctional reinforcement at cell vertices may lead to downstream actomyosin polymerization and contractility. In Drosophila, recycling pathways have well defined roles in cell contact reinforcement, thus Rab25 may be regulating the actomyosin cytoskeleton during epiboly in a similar fashion. Studies in Xenopus have shown that Rab11 recruitment to vertices is modulated by tension (Ossipova et al., 2014). In a positive feedback loop, Rab11 then further promotes actomyosin contractility at these sites to drive apical constriction (Ossipova et al., 2014). Given that tagged-Rab25 localized to vertices, regions under high amounts of tension during cell rearrangements (Bosveld and Bellaïche, 2020), Rab25 may similarly redistribute in response to tension, with the downstream effect of promoting contractility and actin polymerization. Future endeavours will help determine this model of Rab25 function during epiboly.

We propose a two branched model by which Rab25 functions during epithelial epiboly in the zebrafish gastrula. Rab25 vesicle recycling coordinates cytokinetic abscission to ensure successful EVL cell divisions, which is critical for proper cell size, shape and number. In parallel, Rab25 regulates cortical actomyosin network density in the EVL. Increases in cell size from failed cytokinesis led to sparse cortical actin density, thus both Rab25 pathways impact cortical cytoskeleton density in the EVL. Maintenance of the actin cytoskeleton is required for orderly cell rearrangements, polarized force patterns and viscoelastic responses to ensure normal epiboly progression.

## Materials and Methods

### Key Resources Table

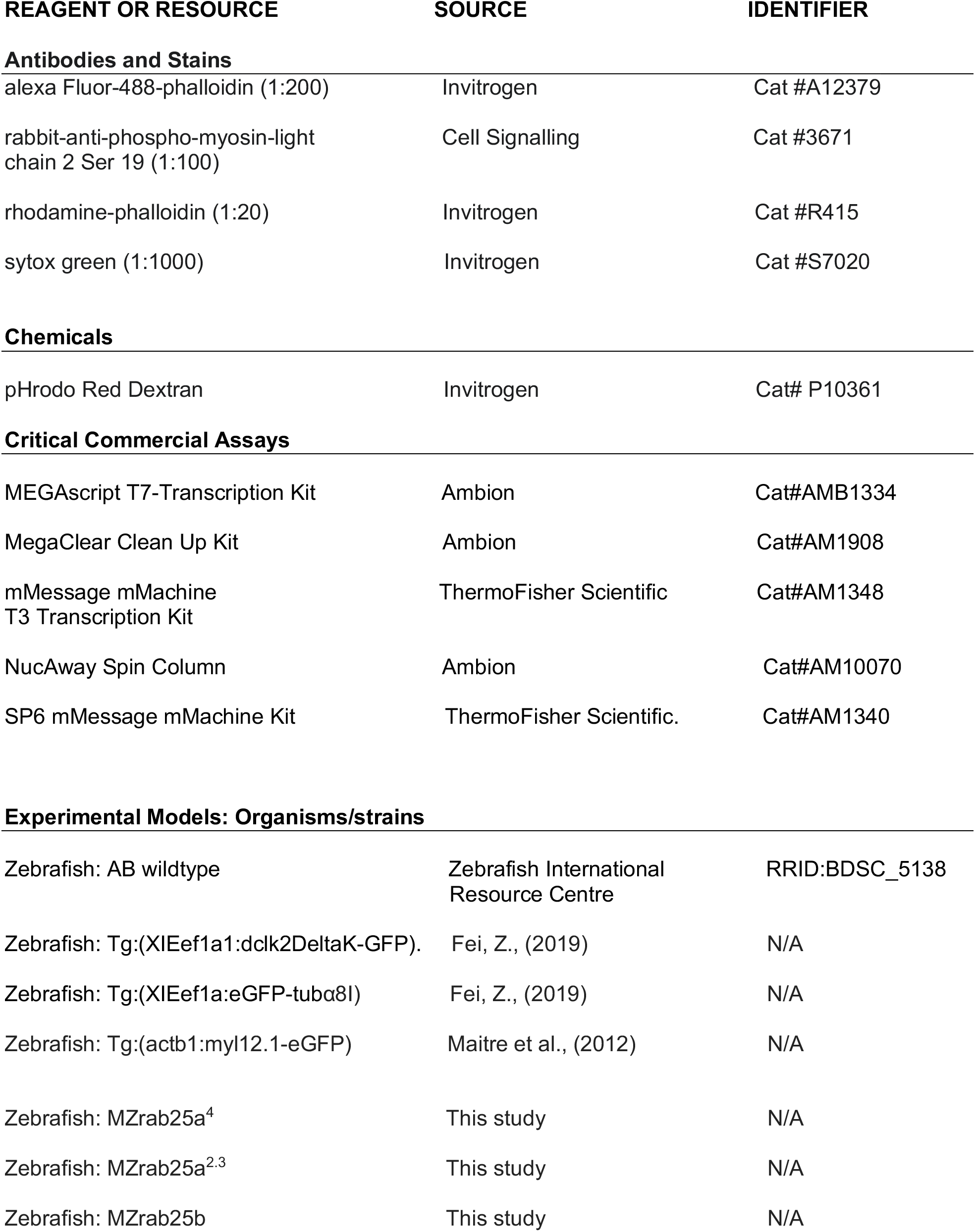
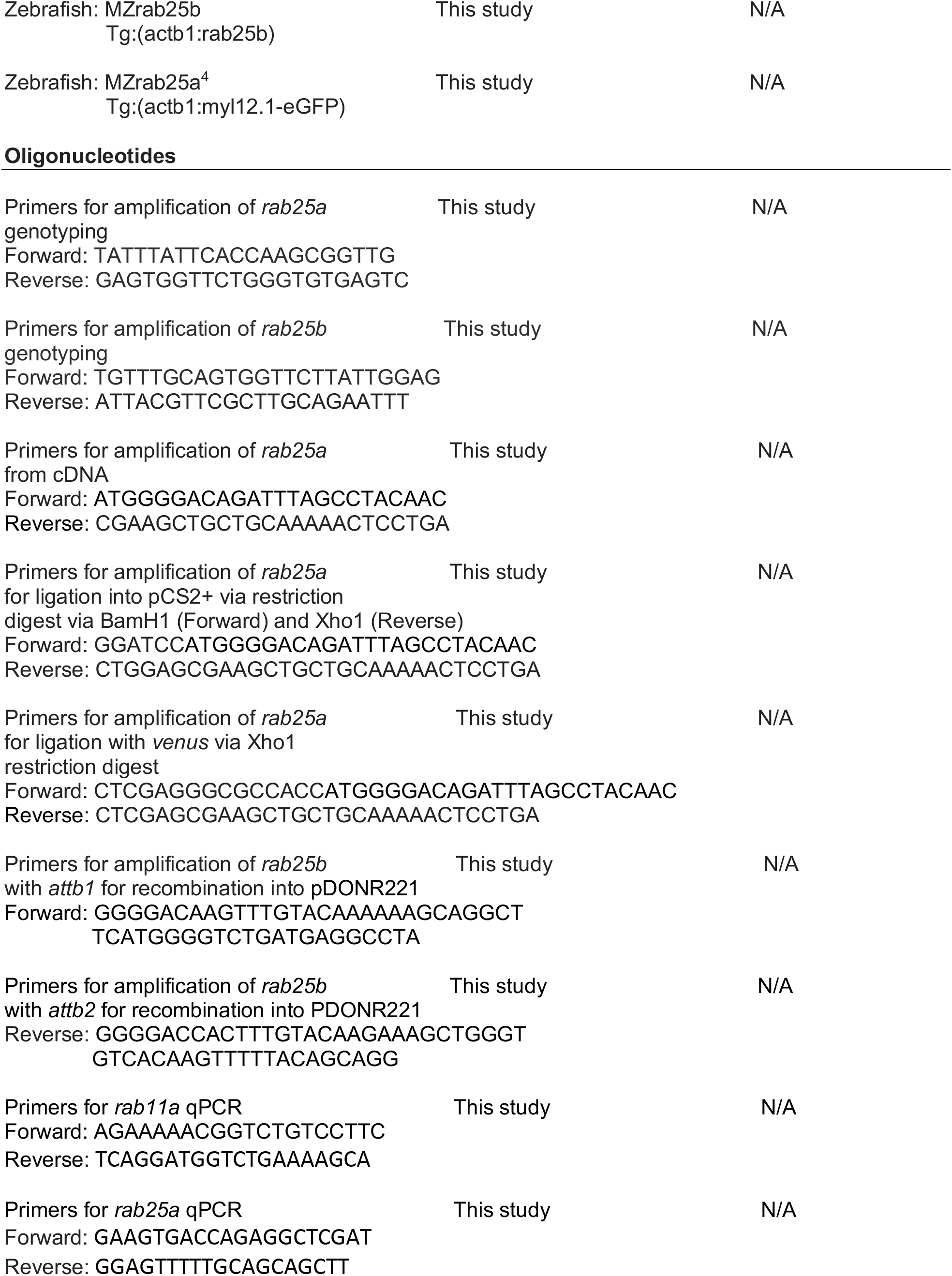
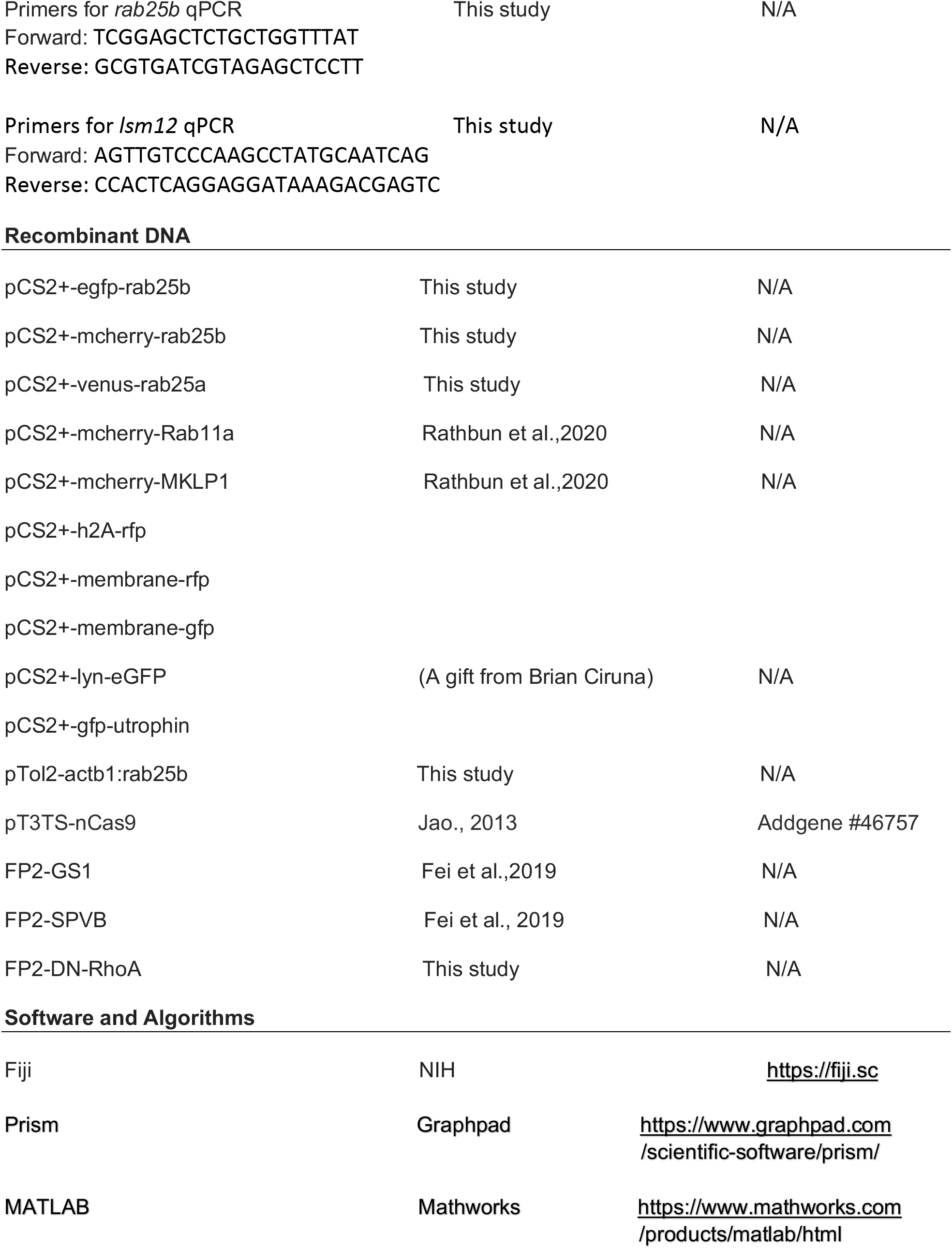
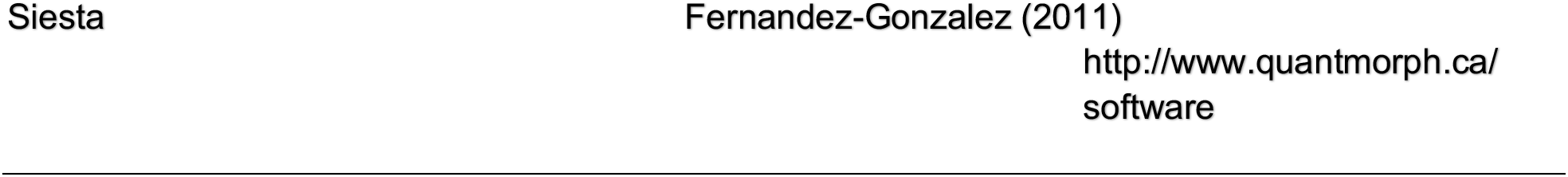

### Zebrafish Handling

Animals were maintained under standard conditions in accordance with the policies and procedures of the University of Toronto animal care committee. Fish stocks were housed at 28-29°C in an Aquaneering Zebrafish Housing System with a pH from 7.2-7.8 and conductivity between 500-600μS. Adults used were on average 1 year old with no prior manipulations or apparent health issues. Embryos were collected from natural spawnings using a tea strainer and rinsed and stored in facility water or E3 medium at 28.5-30°C. Embryos were staged as described by Kimmel et al. (1995).

### Quantification and Statistical Analysis

Quantifications represent data collected from cells visualized within a given area illuminated by confocal microscopy and not estimates of embryo wide measurements.

#### Epiboly Progression

To assess the extent of epiboly progression, the leading edge of the blastoderm was labeled using either phalloidin or by in situ hybridization against ta *(ntla).* Embryos were photographed and ImageJ was used to measure the distance from the animal pole to EVL leading edge and compared to the overall length of the embryo along the animal-vegetal axis (AV) (SF 1D).

#### Cell Contact Analysis

The number of bicellular contacts between an EVL cell and its neighbours EVL cells or the YC was counted.

#### Apical Surface Area, Circularity and Heat Maps

Surface area, circularity and heat maps were calculated and generated using SIESTA (see STAR methods) and custom scripts written in MATLAB.

Circularity=p^2^/4πα,

Where p is the cell apical surface perimeter and α is cell apical surface area. Circularity is 1 for circles and larger than 1 for non-circular shapes.

#### Phalloidin Fluorescence Intensity

Phalloidin levels were measured in fixed embryos in EVL marginal regions at the stages described using a 2μM line at bicellular and tricellular contacts. For each experiment, all measurements were then corrected and normalized to ZO-1 levels in the same regions using the measure tool in ImageJ.

#### Myosin Intensity

Myosin levels were measured across a 2μM line in the middle of bicellular junctions in live embryos expressing myl12.1-eGFP pre-ablation. For each experiment, all measurements were then corrected and normalized to myosin intensity at tricellular contacts along the same junction using the measure tool in ImageJ.

#### pHrodo dextran and lyn-eGFP vesicle counts and fluorescence intensity

In each experiment, vesicles >0.75μM were counted in images from live confocal z-projections of EVL cells. To measure the fluorescence intensity of pHrodo positive vesicles, intensity across a line spanning the diameter of the vesicles was measured. Analysis was done using the measure tool in ImageJ.

#### Cytokinetic Bridge Length and Duration

Bridge length and duration in wildtype embryos was quantified in marginal and animal regions in live AB embryos expressing dcx-gfp from plasmid injection; Tg:(XIEef1a:eGFP-tubα8I) or; Tg:(XIEef1a1:dclk2DeltaK-GFP). All methods produced similar bridge lengths and durations. Mutants embryos expressing dcx-gfp from plasmid injection were used for bridge analysis. Bridge length for all genotypes were measured following intercellular bridge formation. Bridge duration analysis began following cleavage, when microtubules were first organized into bridges. Duration analysis ended when microtubules were no longer visible, indicating abscission. Duration analysis also ended in mutants if bridges were present at the end of time-lapses or when bridges regressed or were torn open. Initial bridge length following formation was measured using the measure tool on ImageJ.

#### Spindle Orientation

The angle of EVL mitotic spindles was compared to the long axis of the cell in live wildtype and mutant embryos in marginal and animal regions using the measure tool in ImageJ. Microtubules in wildtype embryos were labelled by dcx-gfp from plasmid injection; Tg:(XIEef1a:eGFP-tubα8I) or; Tg:(XIEef1a1:dclk2DeltaK-GFP). In mutant embryos, microtubules were labelled with dcx-gfp from plasmid injection. Embryos also expressed membrane-RFP and H2A-RFP.

#### EVL-YC Contact Shortening

The length of EVL-yolk cell contact was measured each frame over the duration of time-lapses using the measure tool in ImageJ.

#### Multicellular EVL-YC vertex resolution

Following formation of multicellular vertices along the EVL-YC boundary, resolution of rosettes was determined when a new cell contact was observed to form between cells adjacent to intercalating cells.

#### Statistics

Supplementary Table 1.

### CRISPR/Cas9 mutant generation

To determine Cas9 target sites against *rab25a* and *rab25b* (ZFIN ID: ZDB-Gene-041212-69; ZDB-Gene-050706-113), the *CHOPCHOP* tool was used (https://chopchop-cbu-uib-no.myaccess.library.utoronto.ca). The following target sites were chosen for *rab25a:* Exon 2 (Rab GTPase Domain), 5’-AGTGGTTTTAATTGGAGAATCAGG-3’; for *rab25b:* Exon 2 (Rab GTPAse Domain), 5’-CTGGATTGGAGCGGTACCGC-3’. To generate gRNA’s, sgDNA templates were generated using PCR without cloning:

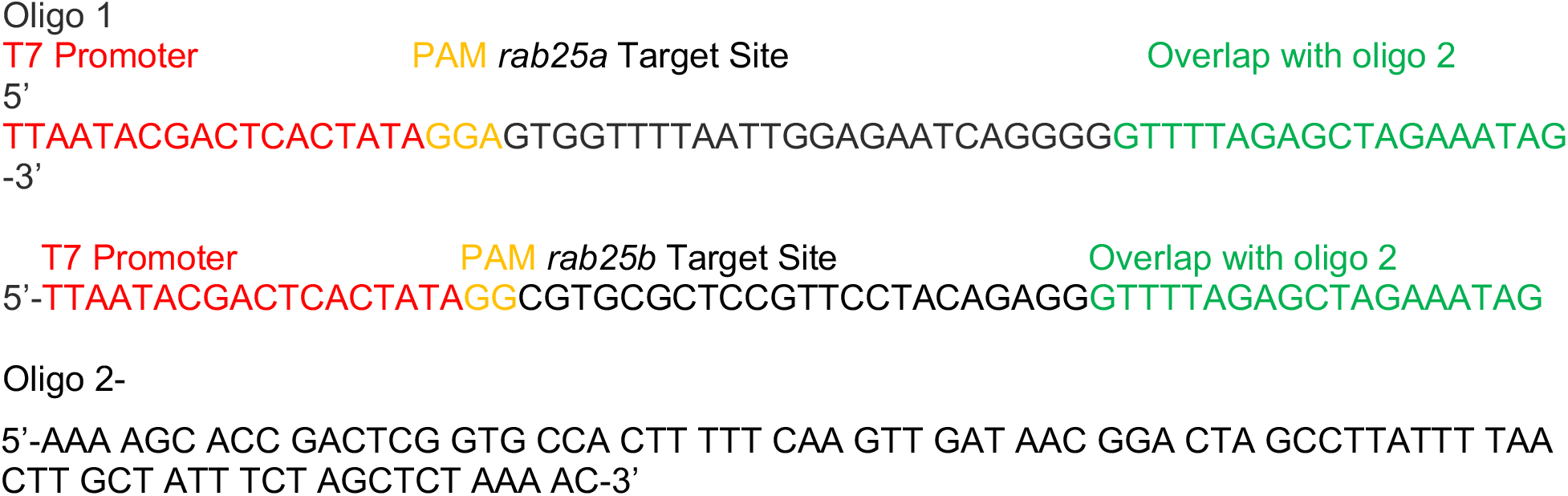

The Ambion MegaScript T7 kit was used to transcribe sgRNA *in vitro.* gRNA (50pg) was coinjected with *cas9* mRNA (300 pg)(pT3TS-nCas9 [Xba1 digest] Addgene plasmid: #46757 Jao., 2013; transcribed using mMESSAGE mMachine T3 kit Life technologies [AM1348])

To determine indel frequencies, genomic DNA from 24hpf injected embryos was extracted using Lysis Buffer and Phenol Chloroform purification with the following primer pairs used for amplification target sites:

*rab25a*

Forward-5’-TATTTATTCACCAAGCGGTTG-3’

Reverse-5’-GAGTGGTTCTGGGTGTGAGTC-3’

*rab25b*

Forward-5’-TGTTTGCAGTGGTTCTTATTGGAG-3’

Reverse-5’-ATTACGTTCGCTTGCAGAATTT-3’

*rab25a and rab25b* PCR products were digested with Hinf1 and Kpn1, respectively. PCR products which could not be cut were then sequenced using the respective forward primers above. This led to the identification of two *rab25a* mutations from two different founder fish and one *rab25b* mutation:

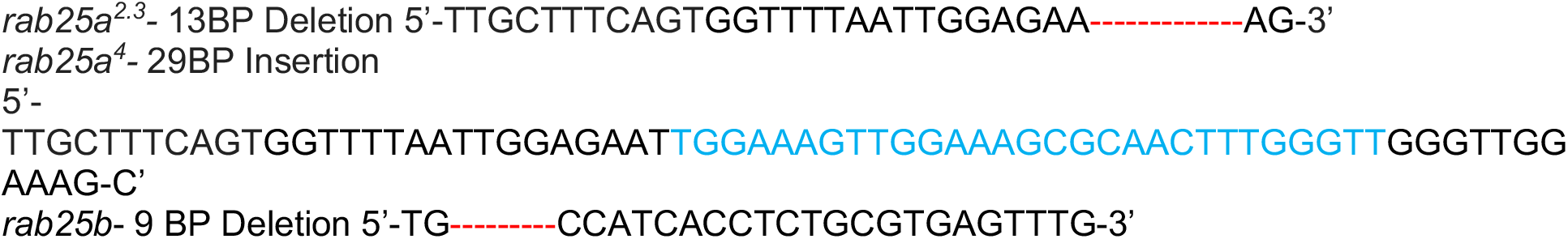

Egg sizes between MZrab25a/b and WT embryos were similar (SF 1G)

We had difficulty rescuing phenotypes through RNA injection, likely due to the required maternal contribution of *rab25a* and *rab25b.* We also observed that *rab25a* mutant alleles produced more severe phenotypes over successive generations of in-crossing.

### Cloning

AB, *MZrab25a* or *MZrab25b* embryos at 24hpf were dechorionated and RNA was extracted from 60-80 embryos using Trizol (Invitrogen). cDNA was generated using Postscript II 1^st^ strand cDNA synthesis kit (NEB). Coding sequences were PCR amplified from cDNA or vectors with Q5 high-fidelity Taq Polymerase (NEB) using the following primers:

*rab25a:* pGEM

Forward: 5’ATGGGGACAGATTTAGCCTACAAC-3’

Reverse: 5’CGAAGCTGCTGCAAAAACTCCTGA-3’

PCR products were gel purified and A-tailed by incubation with dATP and Taq Polymerase (NEB) for 30 minutes and ligated into pGEM-T-Easy (Promega).

*rab25a: pCS2+*

*rab25a* was PCR amplified from pGEM with PCR primers containing Cla1/Stu1 restriction sequences for forward and reverse primers, respectively. PCR fragments were digested with either Cla/Stu1, gel extracted and ligated into pCS2+. Constructs were confirmed by sequencing.

*rab25a: venus-pCS2+*

Forward: 5’-CTCGAGGGCGCCACCATGGGGACAGATTTAGCCTACAAC-3’

Reverse: 5’-CTCGAGCGAAGCTGCTGCAAAAACTCCTGA-3’

N-terminal venus fusion protein was generated by digesting vector and insert with Xho1 and ligated. Sequencing was used to validate ligation of venus and *rab25a.*

*rab25b:*

Forward: 5’-GGGGACAAGTTTGTACAAAAAAGCAGGCTTCATGGGGTCTGATGAGGCCTA-3’ Reverse: 5’-GGGGACCACTTTGTACAAGAAAGCTGGGTGTCACAAGTTTTTACAGCAGG-3’

*rab25b* PCR products were gel extracted and recombined with pDONR221 using BP clonase. Clones were validated by sequencing. Fusion proteins and transgenic vectors were generated by gateway recombination using LR Clonase.

### mRNA synthesis and microinjections

Unless specified otherwise, Not1 digested plasmids were used as templates for *in vitro* transcription using the SP6 mMessage mMachine Kit (Ambion). mRNA was purified using MEGAclear kit or NucAway Spin Columns (Ambion). RNAs were injected into the yolk cell or blastoderm of one-cell stage embryos, as described (Bruce et al.,2003). Doses of injected RNA were: *h2a-rfp (50pg), mrfp/gfp (50pg), lyn-egfp (50pg), gfp-Utrch (100pg), mcherry-rab11a* (300pg), *egfp-rab25b* (300pg), *venus-rab25a* (300pg), *mcherry-rab25b* (300pg) *mcherry-mklp1* (150-200pg). Plasmids were injected at doses ranging from 10-20pg.

### Whole-mount immunohistochemistry

Antibody staining was performed as previously described (Lepage,.2014). Dilutions were as follows: rhodamine-phalloidin (1:200), anti-E-cadherin (Abcam, 1:1000), anti-ZO-1 (1:500), anti-phosphomyosin-light chain 2 Ser 19 (cell signalling,1:100). Embryos were mounted in either 80% glycerol or 0.05% low-melt agarose. Secondary antibodies used were goat-anti-mouse Alexa 488 (Invitrogen, 1:500) and goat anti-rabbit-Cy3 (Jackson immunoresearch,1:500). Sytox green (Invitrogen) was dilution to 0.5μM in fixative.

### Whole-mount *in situ* hybridization

Whole mount in situ hybridizations were performed as previously described (Jowett and Lettice., 1994). *keratin4,ta,rab25a,rab25b,gsc,sox17* probes were generated by linearizing pBS vectors with Not1 and transcribing using T7 polymerase (ThermoFisher Scientific). Probes were purified using NucAway Spin Columns (Ambion).

### phRODO Dextran Assay

Embryos were injected into the yolk cell at the 1-cell stage with *lyn-egfp* mRNA, dechorionated and bathed in pHRODO red dextran (Invitrogen, P10361) dissolved in E3 medium (100ug/ml). WT and *MZrab25b* embryos were incubated in the same media, while *MZrab25a* embryos were in a separate dish. Treatments were performed in petri dishes and embryos were incubated from high stage until 90% epiboly at 30°C. *MZrab25b* embryos were identified by their phenotype and mounted laterally in 0.05% low-melt agarose on glass bottom dishes (Matek).

### Laser Ablations

Tg:(actb1:myl12.1-eGFP) and *MZrab25a* Tg:(actb1:myl12.1-eGFP) embryos at 60% epiboly were dechorionated and mounted laterally in 0.05% low-melt agarose on glass bottom dishes (Matek). Imaging was done at room temp using a Revolution XD spinning disk confocal microscope (Andor Technology) with an iXon Ultra 897 camera (Andor Technology), a 40X (NA1.35; Olympus) oil-immersion lens and Metamorph software (Molecular Devices). Ablations were performed in marginal regions, 2-3 cell rows back from the EVL-yolk cell margin on junctions parallel and perpendicular to the margin. Junctions were cut using a pulsed Micropoint nitrogen laser (Andor technology) tuned to 365 nm. The tissue was imaged immediately before and after ablation in which 10 laser pulses were delivered. 16-bit z-stacks were acquired every 3 seconds at 3um stacks and projected for analysis.

### qPCR

Total RNA was purified from shield stage of zebrafish embryos using Trizol, and an additional DNase I (Turbo DNaseI from Thermo Fisher Scientific) step was used to remove genomic DNA. RNA was reverse-transcribed with random primers using the high-capacity cDNA synthesis kit (Thermo Fisher Scientific). Gene expression was monitored by quantitative real time PCR (qPCR) using primers that distinguish individual transcripts (Supplemental Table 1). The standard curve method was used to calculate expression levels, with zebrafish genomic DNA used to generate the standard curves. Levels of *lsm12b* RNA were used to normalize expression values; primer sequences are shown in Star Methods. All samples were confirmed not to have DNA contamination by generating a reverse transcriptase negative sample and confirming there was no *lsm12b* amplification. At least three biological replicates were analyzed for each experiment. Significant differences in gene expression between shield stage embryos were determined by t-test.

### Imaging

Imaging was performed on a Leica TCS SPB confocal microscope unless otherwise specified.

## Acknowledgements

We are grateful to Heidi Hehnley for the Rab11 constructs. We thank Curtis Boswell for technical advice. For helpful discussion and comments on the manuscript, we thank Rudi Winklbauer and Tony Harris.

## Multimedia

### Video S2. WT EVL Mitosis

WT embryo injected with *mrfp (magenta), gal4* mRNA and *h2A-rfp(magenta)-dUAS-dcx-gfp (*green) plasmid imaged beginning at dome stage. Scale bar: 20μm.

### Video S3. Cytokinetic bridge regression in *Mrab25b* mutant embryo

*Mrab25b* embryo injected with *mrfp (magenta)* and *gal4* mRNA and *h2A-rfp*(magenta)-dUAS-*dcx-gfp (*green) plasmid imaged beginning at dome stage. Scale bar: 20μm.

### Video S4. Failed cytokinetic midbody formation in *MZrab25b* embryo

(Left panel) Tg:(dcx-gfp) (magenta) expressing mcherry-hklp1 (green) imaged beginning at dome stage showing formation of midbody in WT embryo (right panel). Apical z-projection of a *Mrab25b* embryo expressing mRfp (magenta) and mcherry*-*Mklp1 (green); failed coalescence of the mcherry-Mklp1 precedes bridge regression. Scale bar: 20μm.

### Video S5. Torn open intercellular bridges during multipolar mitosis in *Mrab25b* embryo

*Mrab25b* embryo injected with *mrfp (magenta)* and *gal4* mRNA and *h2A-rfp*(magenta)-dUAS-*dcx-gfp (*green) plasmids imaged beginning at dome stage. Scale bar: 20μm.

### Video S6. EVL cells interconnected by long cytokinetic bridges in *Mrab25b* embryo

*Mrab25b* embryo injected with *mrfp (magenta)* and *gal4* mRNA and *Λ2A-rfp*(magenta)-dUAS-*dcx-gfp (*green) plasmid imaged beginning at dome stage. Scale bar: 20μm.

### Video S7. Heterogenous distribution of F-actin in *MZrab25b* EVL cells during late epiboly

WT embryo expressing Utrch-GFP (Fire-LUT) imaged beginning at 7hpf, showing cortical F-actin enrichment during marginal EVL cell rearrangements (left panel). *MZrab25b* embryo expressing Utrch-GFP (Fire-LUT) imaged beginning at 7hpf (right panel). Cells with reduced and large apices shorten their EVL-YC contact (right panel). Scale bar: 20μm.

### Video S8. venus-Rab25 is recruited to vertices of rearranging marginal cells

WT embryo expressing venus-Rab25a beginning at 7hpf, enrichment can be seen at the marginal tip of the intercalating cell. Scale bar: 20μm.

### Video S9. Myosin-Gfp distribution in *MZrab25b* embryo during epiboly progression

WT Tg(Myl1.1-Gfp) (Fire-LUT) embryo imaged starting at 7hpf (left panel). *MZrab25b* Tg(Myl1.1-Gfp) (Fire-LUT) embryo imaged starting at 7hpf (right panel). Myosin-Gfp becomes heterogeneously distributed in EVL marginal regions. Scale bar: 20μm.

---

**SF1.**
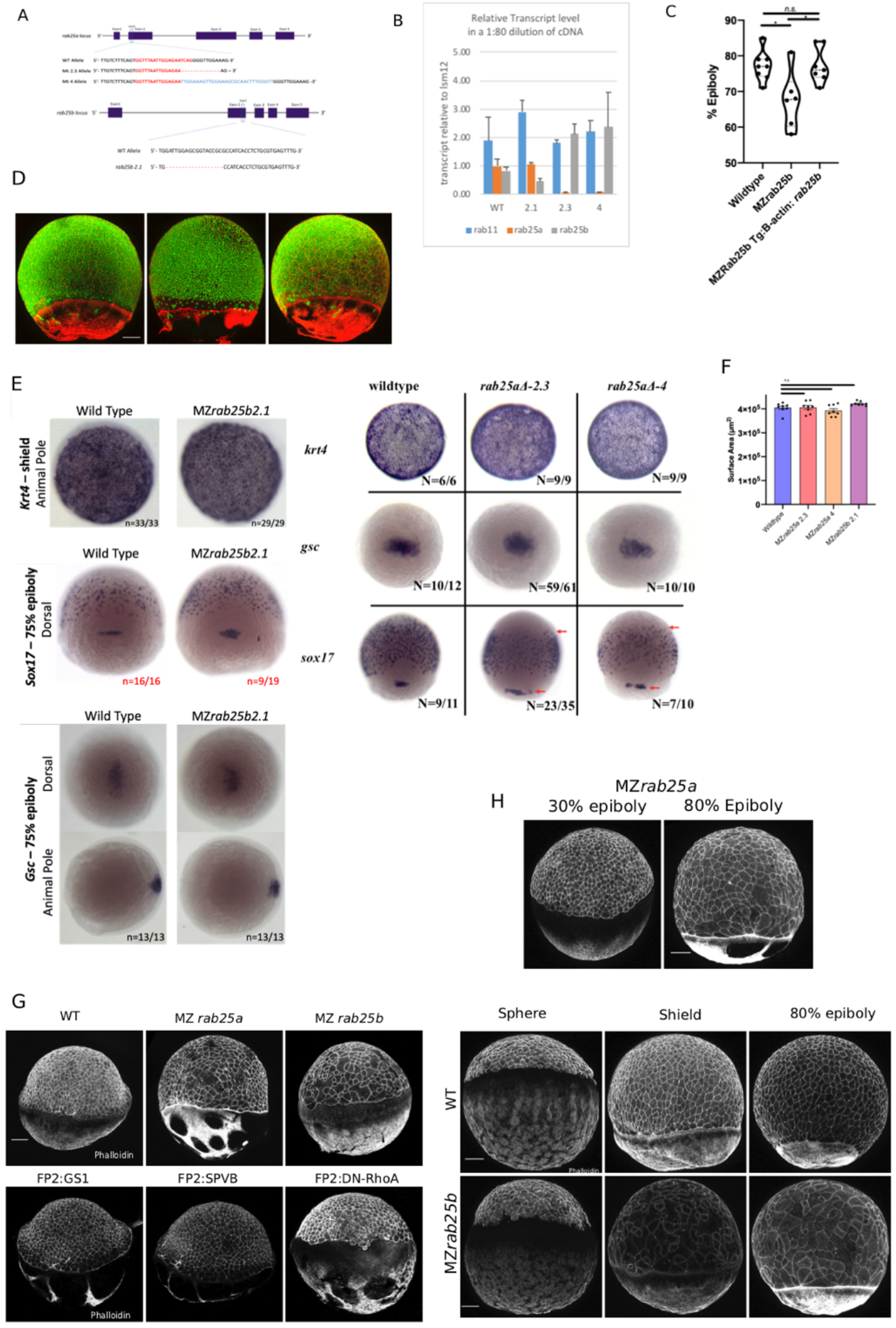
Maternal-Zygotic Rab25a and Rab25b characterization. (A) Schematic of *rab25a* and *rab25b* loci with Cas9 target sites and mutant allele sequences. (B) qPCR analysis of *rab25a, rab25b* and *rab11a* transcripts levels in WT, *MZrab25a* and *MZrab25b* embryos at shield stage. (C) Tg: MZrab25b (B-actin;*rab25b*) epiboly movements rescued compared to MZrab25b mutants. WT (n=8), *MZrab25b* (n=6) and *MZrab25b* Tg (B-actin; *rab25b)* (n=7) time-matched at 8hpf. Means;sem; One-Way ANOVA, * p<0.05. (D) Stage-matched WT, *MZrab25a* and *MZrab25b* embryos at 75% epiboly showing phalloidin staining of f-actin (red) and sytox green labeling of nuclei (green). Embryos positioned laterally with animal pole to the top; Scale bar: 100μm (E) Whole-mount *in situ* hybridizations of indicated probes in purple at indicated stages. Scale bar: 100μm. Arrows indicate sox 17 staining. (F) Surface area of laterally positioned embryos at shield stage. WT (n=8), *MZrab25a* 2.3 (n=8), *MZrab25a* 4 (n=8) and MZr*ab25b* (n=8) embryos sizes at 6hpf. Means; SEM; One-Way ANOVA. (G) Epithelium does not phenocopy mutant embryos when WT yolk cell F-actin networks are perturbed. Stage-matched embryos positioned laterally at 6hpf stained for F-actin (phalloidin). Scale bar: 100μm. (H) Epithelial Defects emerge over the duration of epiboly in Rab25 mutant embryos. Stage-matched WT, *MZrab25a* and *MZrab25b* positioned stained for F-actin (phalloidin). Scale bar: 100μm.

**SF2.**
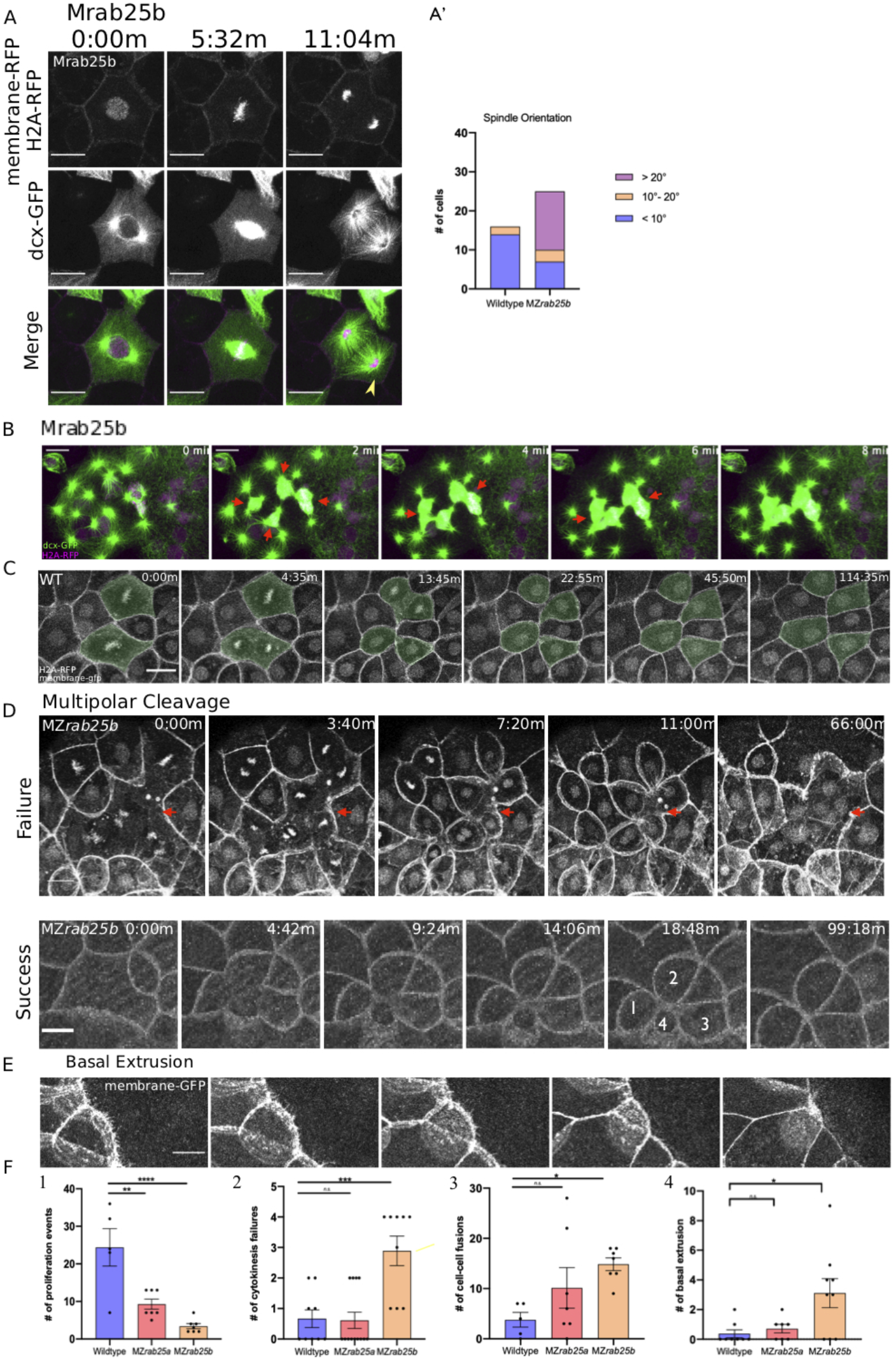
Cytokinesis related defects in MZRab25b embryos. (A) Z-projections of stills from confocal time lapse of *MZrab25b* mononucleate EVL cell during mitosis at 30% epiboly. Microtubules (green), membrane (magenta) and nuclei (magenta). Arrowhead indicates reduced density of astral microtubules. Relative proportions of spindle orientation along the long axis of cells during mitosis. Scale bar: 20μm. (B) Z-projections of stills from confocal time lapse of *MZrab25b* mitotic spindles fusing during multipolar cytokinesis. Time-lapses began at 30% epiboly in marginal regions; arrows denote fusing spindles; Microtubules (green), membrane (magenta) and nuclei (magenta). Scale bar: 10μm. (C) Z-projections of stills from confocal time lapse of a WT embryo imaged beginning at dome stage. Shaded cells are dividing; Nuclei and membranes labelled. Scale bar 20μm. (D) Z-projections of stills from confocal time lapse of multipolar mitosis in MZrab25b embryos beginning at 30% epiboly. Failed cleavage indicated by arrows; Successful cleavage generates cells with reduced apices indicated by numerals; Nuclei and membranes labelled. Scale bar: 10μm. (F) Quantifications of means of (1) proliferation events in WT (n=5), *MZrab25a* (n=7) and *MZrab25b* (n=7);(2) Cytokinesis events in WT (n=9), *MZrab25a* (n=13) and *MZrab25b* (n=9); (3) non-sister cell fusions in WT (n=5), *MZrab25ba* (n=7) and *MZrab25b* (n=7); (4) Basal Cell Extrusion events in WT (n=8), *MZrab25a* (n=7) and *MZrab25b* (n=9). means; SEM; One-Way ANOVA; *,**,***,*o**, p<0.05,0.01,0.001,0.0001.

**SF3.**
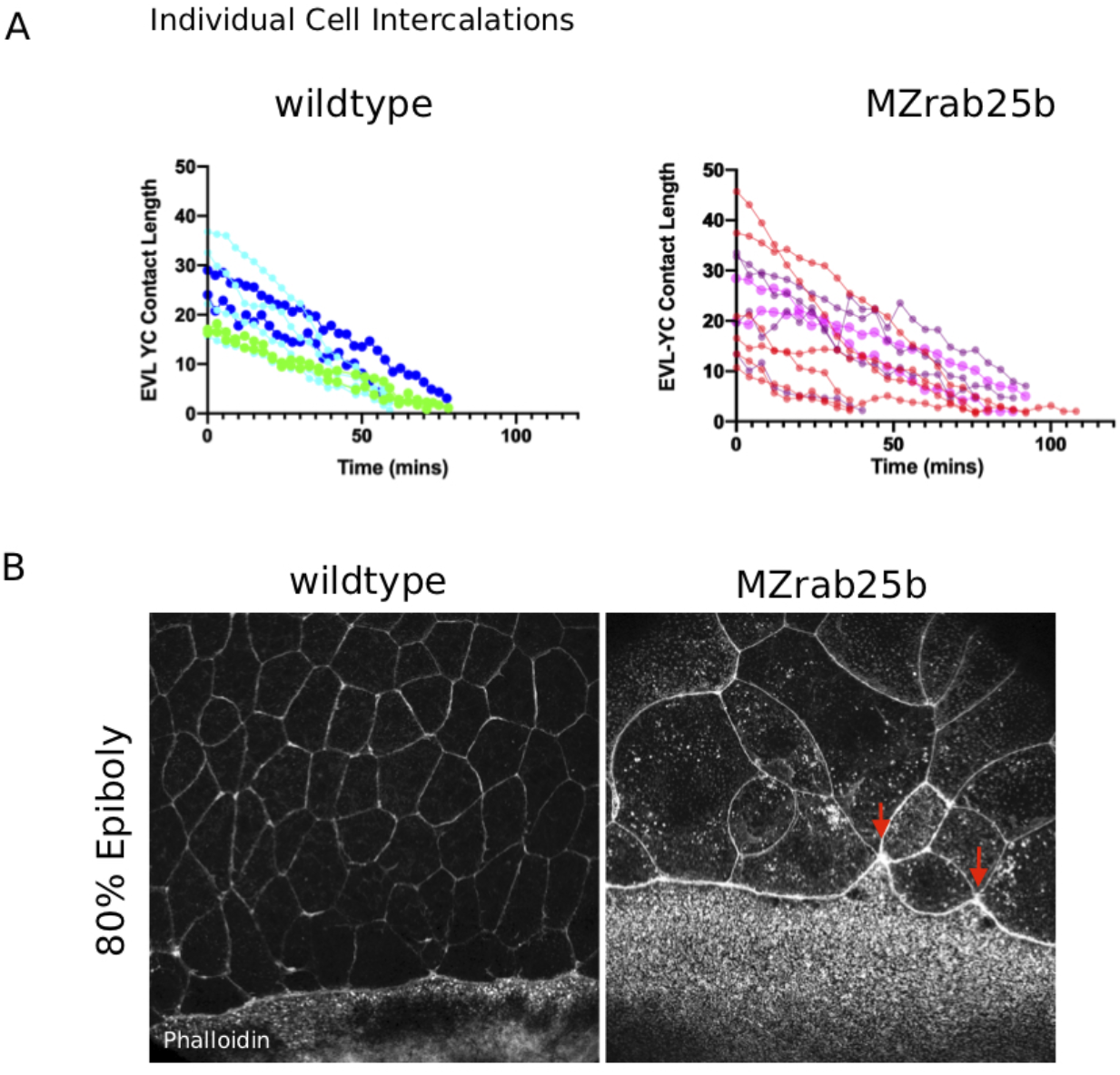
Marginal EVL cell behaviours during epiboly progression. (A) Individual marginal cell EVL-YC contact length over time in WT (n=8, N=3) and *MZrab25b* (n=13, N=3) beginning at 7hpf. (B) Z-projections of stills from confocal time lapse of stage-matched WT and *MZrab25b* embryos during late epiboly stained for F-actin. Arrowhead denotes uneven EVL margin in MZrab25b embryos; Scale bar: 20μm.

**SF4.**
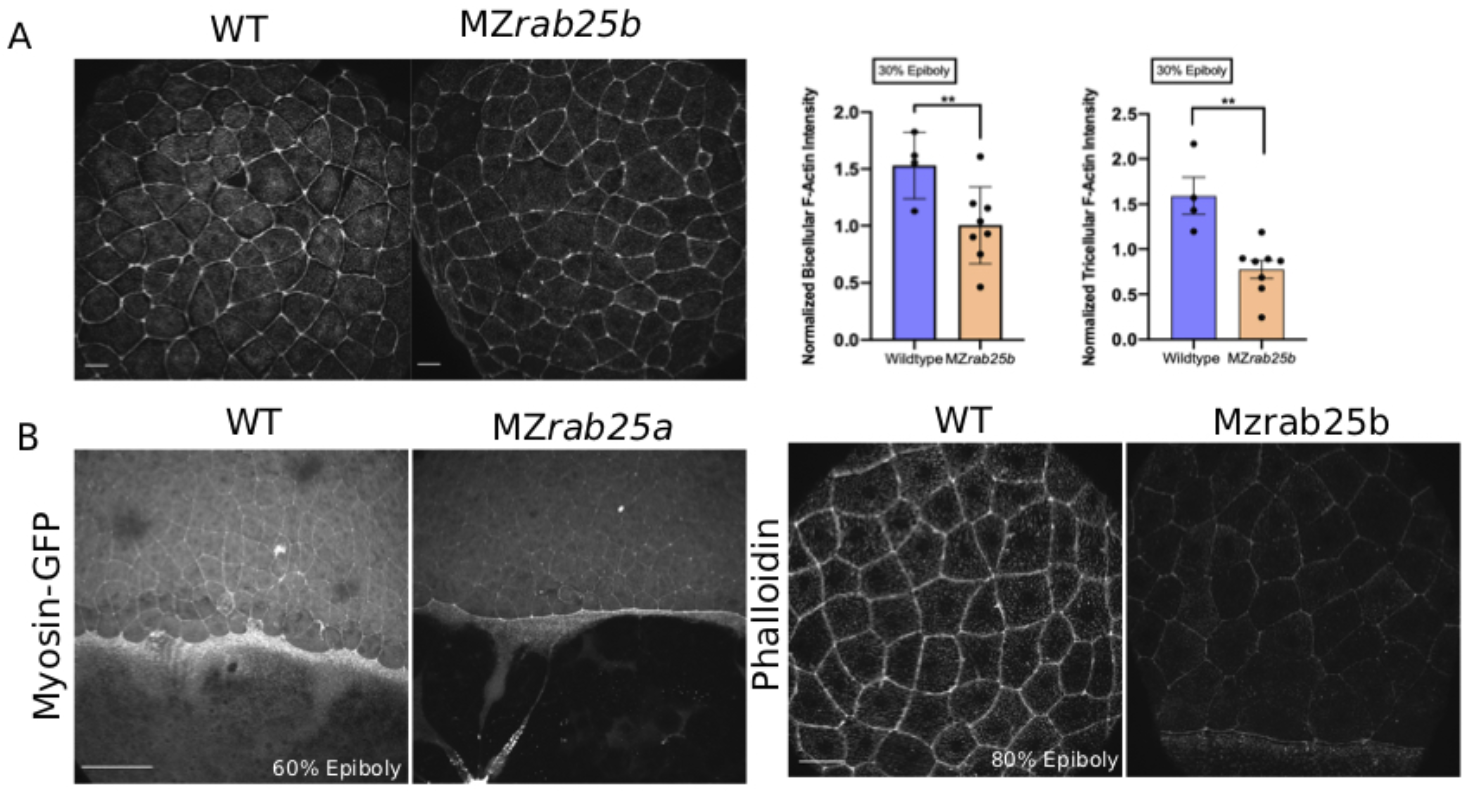
Actomyosin intensity in MZrab25a and MZrab25b epiboly. (A) Confocal z-projections WT and *MZrab25b* embryos stage-matched at 30% epiboly and stained for f-actin (phalloidin). Quantifications of EVL tri-or bicellular f-actin intensity in WT (n=40, N-4) and *MZrab25b* embryos (n=70, N=7). Means;SEM; t-test **,P<0.01. Scale bar 20μm. (B) Confocal z-projection of stage-matched WT Tg(Myl1.1-gfp) and *MZrab25a* Tg(Myl1.1-gfp) at 60% epiboly positioned laterally. Confocal z-projection of stage-matched WT and *MZrab25b* embryos stained for F-actin (phalloidin). Scale bars 20μm.

**SF5.**
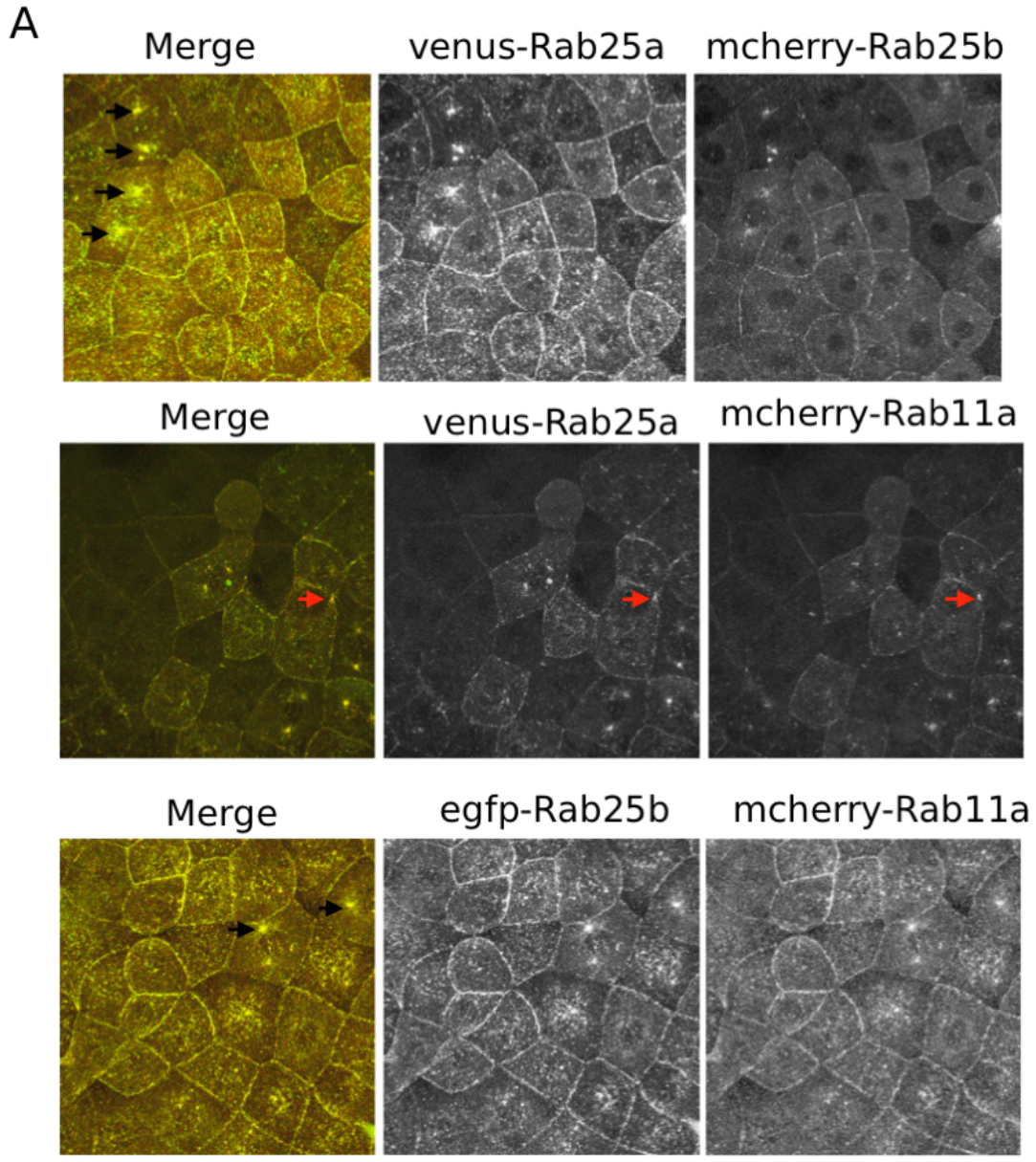
Co-localization of N-terminally fluorescently tagged Rab11a, Rab25a and Rab25b constructs. (A) Confocal z-projections of N-terminally fluorescently tagged Rab constructs at 30% epiboly in WT embryos. Constructs co-localize at the plasma membrane, cytosol, centrosomes (black arrows) and tricellular vertices (red arrows); Scale bar: 20μm.

